# Comparative multi “omics” profiling of *Gossypium hirsutum* and *Gossypium barbadense* fibers at high temporal resolution reveals key differences in polysaccharide composition and associated glycosyltransferases

**DOI:** 10.1101/2025.04.26.650795

**Authors:** Sivakumar Swaminathan, Youngwoo Lee, Corrinne E. Grover, Megan F. DeTemple, Alither S. Mugisha, Lauren E. Sichterman, Pengcheng Yang, Jun Xie, Jonathan F. Wendel, Daniel B. Szymanski, Olga A. Zabotina

## Abstract

Among the two allopolyploid cultivated species of cotton, *Gossypium barbadense* is known for its superior quality fiber compared to *G. hirsutum*. Length and strength are key determinants of the fiber quality. Although the mature fibers are dried cell walls consisting mainly cellulose, the dynamic remodeling of pectin, xyloglucan, and xylan polysaccharides during fiber growth significantly impact the final fiber quality. Comprehensive knowledge about polysaccharides and their biosynthesis during fiber development of the cultivated species is important for improving fiber quality. In this study, comparative large-scale glycome, transcriptome and proteome profiling were conducted daily on fibers of both cotton species covering critical stages of fiber development spanning primary cell wall synthesis and the transition to secondary wall synthesis. Interspecific comparisons revealed that a delayed accumulation of cellulose content, as well as the occurrence of lower levels and differential compositions of non-fucosylated/fucosylated xyloglucans, homogalacturonans, and highly branched Rhamnogalacturonan-I polysaccharides might be contributing to longer fiber phenotypes of *G. barbadense* relative to *G. hirsutum*. Our study also suggests, differential temporal compositions of heteroxylans might contribute to variation in cellulose microfibril arrangement and strength of fiber exists between the two species of cotton. Comparative transcriptomic analysis identified differentially expressed polysaccharide-synthesizing glycosyltransferases that might underlie fiber quality differences between the two species. Transcripts encoding many cell wall localized expansins were more abundant in *G. barbadense* than in *G. hirsutum*. Overall, these findings extend our knowledge regarding the molecular factors that contribute to fiber quality and provide insights for targeted cotton fiber improvement.

**SIGNIFICANCE STATEMENT:** Comparative multi-omics profiling of two major commercial cotton species, *Gossypium hirsutum* and *Gossypium barbadense* revealed substantial differences in polysaccharide structures and expressed polysaccharide-synthesizing glycosyltransferases, that potentially contribute to differences in fiber length and strength. The molecular details elucidated in the present study contribute to the goal of improving fiber quality and its commercial value.

## INTRODUCTION

Worldwide, cotton (*Gossypium* spp.) is the most important natural fiber used in the textile industry and it is cultivated in over 80 countries. Among 50 naturally occurring cotton species, four have been domesticated, namely, two allotetraploids from Central and South America, *Gossypium hirsutum* (Upland or American cotton; AD1 genome), and *G. barbadense* (Egyptian or Pima cotton; AD2 genome), respectively, and two African-Asiatic diploids *G. arboreum* (Tree cotton; diploid A2 genome) and *G. herbaceum* (Levant cotton; diploid A1 genome) (Hu et al., 2021; Jareczek et al., 2023; Viot & Wendel, 2023). About 98% of the commercially cultivated cotton worldwide is from *G. hirsutum* (*Gh*) and *G. barbadense* (*Gb*). *Gh* offers higher yield and wider environmental adaptability, but only moderate fiber quality suited for general-purpose textiles. In contrast, *Gb* cotton is grown only in selected environments and its fiber is longer, stronger, thinner and finer (having less mass per unit length), which makes it preferred for spinning softer yarns used for luxury cotton clothing.

Cotton fibers are single cells that elongate from the epidermis of the cotton seed coat, beginning near the day of anthesis (DPA) and reaching about 2.5 to 4 cm or more in length during 50 to 55 DPA. Cotton fiber development is a highly coordinated, modular process consisting of at least five overlapping stages: 1) fiber cell initiation and tapering (–3 to 3 DPA); 2) elongation stage of primary cell wall (PCW) development (2 to 3 weeks); 3) transition period of PCW remodeling/early secondary cell wall (SCW) formation; 4) high cellulose accumulation/SCW thickening (3 to 5 weeks); and 5) maturation/drying (6 to 8 weeks) (Haigler et al., 2012; Kim, 2015; Huang et al., 2021; Yanagisawa et al., 2022). On an average, longer fibers develop in *Gb*, perhaps due to its extended elongation phase in comparison to *Gh* (Hu et al., 2019).

The spinnable cotton fiber cell is primarily composed of dried cell wall (CW) polysaccharides. Cotton fiber development involves synchronized gene expression networks, hormone signaling and biosynthetic pathways, and physiological and developmental processes that drive dynamic changes in fiber CW polysaccharide composition (Jan et al., 2022; Jareczek et al., 2023; Swaminathan et al., 2024; Grover et al., 2025). The initial PCW is rich in pectic polysaccharides and xyloglucans (Huwyler et al., 1979; Maltby et al., 1979; Singh et al., 2009), which confer plasticity to the CW, enabling rapid fiber elongation under the influence of high internal turgor pressure. As the fiber transitions to SCW production, pectins and xyloglucans decrease in relative abundance and, in combination with the increase of cellulose content, heighten CW rigidity and strength (Tokumoto et al., 2002). After the transition stage, SCW entails massive cellulose accumulation in fibers, followed by maturation, programmed cell death, lumen collapse, and fiber dehydration. The matured, dried fibers used for yarn and textile manufacturing contain more than 95% cellulose with traces of non-cellulosic polysaccharides, glycoproteins, sugars, and minerals (Haigler et al., 2012).

The availability of superior quality *Gb* and moderate quality *Gh* accessions and temporally sampled polysaccharides during fiber PCW and SCW synthesis make the cotton fiber an excellent model to gain insight into the molecular and biochemical processes responsible for cotton fiber development (Lacape et al., 2012; Li et al., 2013; Tuttle et al. 2015; Hu et al., 2019; Zhang et al., 2022; Liu et al., 2023). This understanding, in turn, will inform efforts to improve fiber quality traits in both allopolyploid cotton species.

Comparative “glycomic” studies on fiber CW polysaccharides across different cotton species have established that dynamic changes occur in the non-cellulosic polysaccharide epitopes of pectins/xyloglucans/xylans, in addition to cellulose, during fiber elongation, transition, and early SCW thickening stages (Singh et al., 2009; Avci et al., 2013; Hernandez-Gomez et al., 2015; Hernandez-Gomez et al., 2017; Guo et al., 2019; Pettolino et al., 2022; Swaminathan et al., 2024). These studies show that fiber length in various species is determined during the PCW biosynthesis/remodeling in the elongation stage, while fiber strength is influenced during the SCW biosynthesis/thickening stage. In addition to CW polysaccharide composition and the extended fiber elongation stage, subtle variations in the timing of cellulose deposition in fiber CW have been implicated as an important factor for superior quality (long, strong and finer) of *Gb* fiber in comparison to *Gh* (Avci et al., 2013; Li et al., 2013; Rajasundaram et al., 2014).

Earlier, we carried out glycome, transcriptome, and proteome profiling on *Gh* (accession TM-1) and identified critical polysaccharide epitopes and putative potential glycosyl transferases (GTs) that synthesize various polysaccharide epitopes (Swaminathan et al., 2024; Grover et al., 2025; Lee et al., 2025). In the present study, we studied *Gb* (accession 3-79) using precisely the same approaches and compared the two-cotton species (*Gh* and *Gb*) with respect to glycome, and transcriptome profiling using temporally dense sampling, in parallel to our earlier study (Swaminathan et al., 2024). Thus, we compared developing fibers of *G. hirsutum* (accession TM-1) and *G. barbadense* (accession 3-79) daily from 6 to 25 DPA. These 20 consecutive days span PCW into SCW synthesis, where gene expression variation between *Gh* and *Gb* are different (Al-Ghazi et al., 2009; Liu et al., 2023). A broad collection of polysaccharide-specific monoclonal antibodies was utilized for glycome profiling. Further, a comparative analysis between the polysaccharide epitopes and glycosyltransferase genes (from transcriptome data) involved in synthesizing the polysaccharide epitopes was carried out. Although the CW polysaccharides epitope content during different stages of fiber development is changed by the presence of both the glycosyltransferases and CW glycosyl hydrolases, in this study, we focused exclusively on the glycosyltransferases. Our large-scale, high temporal resolution comparative multi-omics analyses between two cotton species revealed many dynamically remodeled polysaccharides and the associated differentially expressed glycosyltransferases that might contribute to the different fiber quality traits in the two species. These findings provide insights that could inform cotton breeding strategies for improved fiber traits.

## RESULTS

### Polysaccharide composition of *G. hirsutum* (*Gh*) and *G. barbadense* (*Gb*) cotton fibers during development (6 - 25 DPA)

The buffer-soluble fractions (50mM CDTA:50mM ammonium oxalate buffer extracts), alkali-soluble fractions (4M KOH extracts), and cellulose were extracted from cotton fiber CW (6 to 25 DPA), lyophilized, weighed and summarized (Figure 1; Table S1). As expected, the amount of cellulose and non-cellulosic polysaccharide fractions per boll increased gradually from 6 to 25 DPA in both species.

**Figure 1.**
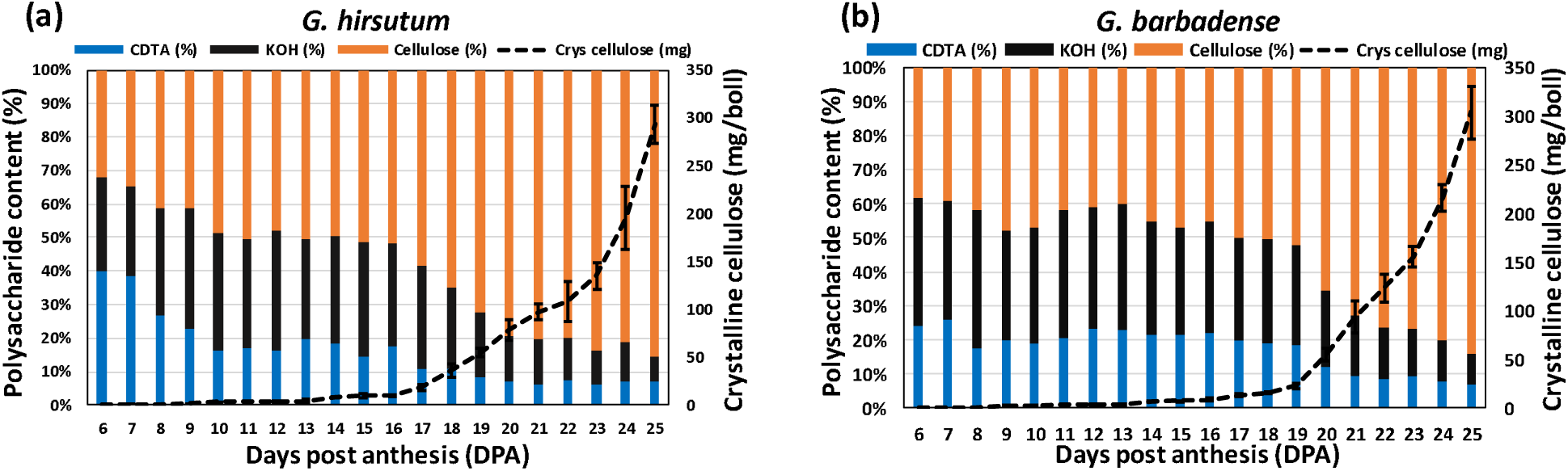
Polysaccharide contents of fiber cell wall (CW) fractions in G. hirsutum (Gh) and G. barbadense (Gb) during development (6 - 25 DPA). (a) Content of buffer-soluble (50mM CDTA-50mM Ammonium oxalate extract), alkali-soluble (4M KOH extract), total cellulose, and crystalline cellulose polysaccharide fractions in G. hirsutum fibers. Data represent averages from three biological replicates. (b) Content of buffer-soluble (50mM CDTA-50mM Ammonium oxalate extract), alkali-soluble (4M KOH extract), total cellulose, and crystalline cellulose polysaccharide fractions in G. barbadense fibers. Data represent averages from three biological replicates.

At 6 DPA, total cellulose content (consists of both amorphous and crystalline cellulose) was 33.6% and 38.4%, for *Gh* and *Gb*, respectively (Figure 1; Table S1). Cellulose content gradually increased in both species after this time; however, the rate of increase was slower in *Gb*. By 16

DPA, cellulose content reached 51.6% in *Gh* and 45.4% in *Gb*. After 16 DPA, the cellulose content of *Gh* started to increase rapidly (61.9% at 17 DPA; 72.9% at 19 DPA), coinciding with the SCW thickening stage (Li et al., 2013) and reached 85.5% at the end of 25 DPA. Interestingly, unlike *Gh*, the cellulose content of *Gb* increased gradually and slowly after 16 DPA and reached only about 51.9% at the end of 19 DPA, and after that, it started to increase rapidly (65.4% at 20 DPA; 72.8% at 21 DPA), coinciding with the SCW thickening stage, reaching 84.1% at the end of 25 DPA, a comparable level to that of *Gh*. The amount of estimated crystalline cellulose content of both species showed profile patterns similar to that of total cellulose content. Crystalline cellulose content was about 0.2 mg per boll at 6 DPA and about 9.0 mg at 16 DPA in both species (Figure 1; Table S1). From 17 DPA onwards, crystalline cellulose content increased rapidly in *Gh*, but at slower rate in *Gb*. At 20 DPA, crystalline cellulose content 78.2 mg in *Gh* and 54.7 mg in *Gb*. After 20 DPA this amount increased rapidly (at 21 DPA, 97.8 mg in *Gh* and 92.8 mg in *Gb*). At 25 DPA, the crystalline cellulose content was estimated to be 293.6 mg in *Gh* and 303.5 mg in *Gb*. Overall, in *Gb* there is a delay of ∼3 days in cellulose content accumulation relative to *Gh*.

The content of buffer-soluble polysaccharides at 6 DPA was significantly lower in *Gb* (24.3%) in comparison with *Gh* (39.4%). Relative content of the buffer soluble polysaccharide fraction gradually decreased after 6 DPA and the reduction was faster after 17 DPA for *Gh* and 20 DPA for *Gb*, which coincided with their respective increase in cellulose content. The relative amount of buffer soluble polysaccharide fractions reached 6.9% in *Gh* and 7.0% in *Gb* at the end of 25 DPA. The content of alkali-soluble polysaccharides was 27.0% for *Gh* and 37.3% for *Gb* at 6 DPA and 30.6% and 32.4%, respectively, at the end of 16 DPA. Similar to the buffer-soluble polysaccharide fraction, the relative content of alkali-soluble polysaccharides started to decrease at a faster rate after 16 DPA, coinciding with an increase in the cellulose content, and reached 7.6% and 8.9% in *Gh* and *Gb*, respectively at the end of 25 DPA.

#### Glycome, transcriptome, and proteome analyses of *Gb* fiber

We performed glycome, transcriptome, and proteome profiling of *Gb* fibers sampled each day between 6 and 25 DPA. As described in our previous study (Swaminathan et al., 2024), glycome profiling of cotton fiber CW polysaccharide epitopes were performed using 71 different monoclonal antibodies (Figure S1; Table S2) and the glycome profiling data of *Gb* fiber presented in Table S3. To identify the polysaccharide-synthesizing glycosyltransferases (Table S4) that contribute to the epitope patterns observed in *Gb* glycome profiling (Table S3), we analyzed the transcriptome profiling data (deposited in NCBI-SRA under PRJNA1222456) and the proteome data collected from the microsomal fraction (P200) (Table S5) prepared at each DPA (6 to 25 DPA). Here, we focus on polysaccharide-synthesizing glycosyltransferases and associated polysaccharide-decorating enzymes (methyl- and acetyltransferases) responsible for synthesizing the polysaccharides that are analyzed in the glycome profiles. From the total of 18,844 detected P200 proteins, we identified 93 glycosyltransferases (Table S3). Using glycome, transcriptome, and proteome profiling data obtained from *Gb* fibers, we carried out Pearson correlation coefficient (PCC) analysis and identified significantly correlated profiles of polysaccharide epitopes and corresponding glycosyltransferases from both transcriptome and proteome (Table S6) (representative correlated profiles shown in Figure 2). For example, four xyloglucan xylosyltransferases (XXTs) (GbXXT2-1Dt, GbXXT2-2At, GbXXT3-1At, GbXXT3-2At), involved in xyloglucan biosynthesis, detected in the proteome showed protein expression profiles similar to their corresponding transcripts and xylosylated xyloglucan epitopes (GbXG-XX/GbXG-XXXG), with levels gradually decreasing from 6 to 25 DPA (Figure 2a; Table S6). Similarly, the pattern of three xyloglucan galactosyltransferases (GbMUR3-2At, GbMUR3-2Dt, GbMUR3-4Dt) from proteome were found to correlate with levels of the transcripts and the galactosylated xyloglucan epitopes (GbXG-XLX/GbXG-L) (Figure 2b; Table S6). One of the fucosyltransferase (GbFUT1-2At) from both transcriptome and proteome was significantly correlated to the three of the fucosylated xyloglucan epitopes (Table S6).

**Figure 2.**
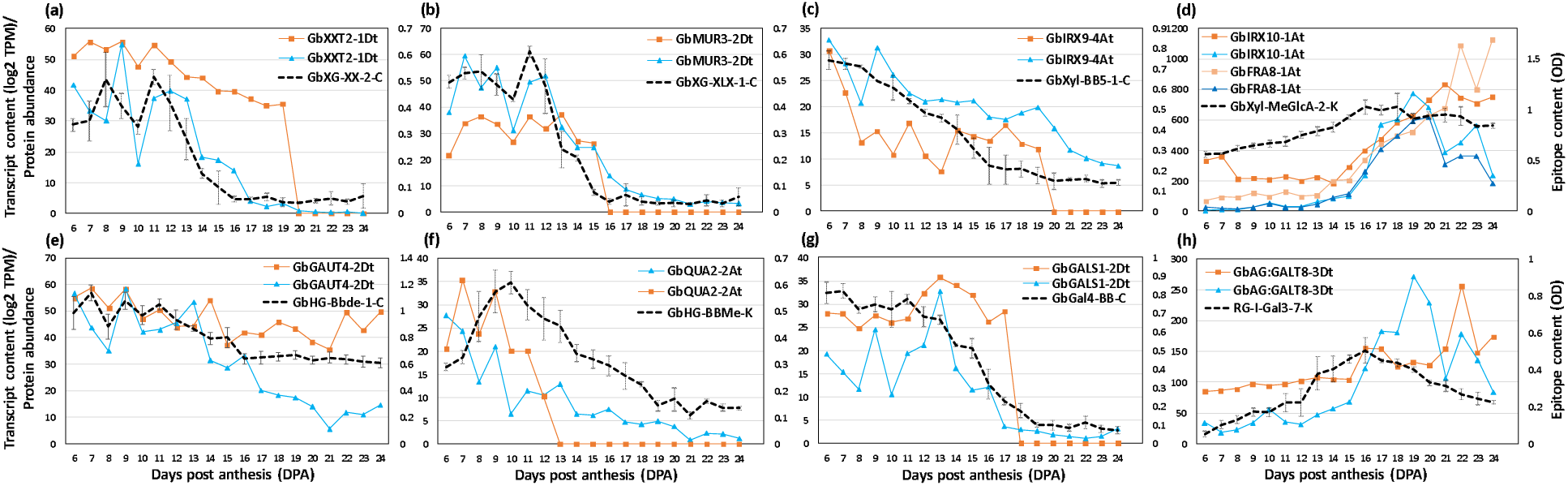
Representative profiles of G. barbadense polysaccharide epitopes and significantly correlated transcripts and protein profiles of glycosyltransferases involved in synthesizing these epitopes. Polysaccharide epitopes are represented by a black dotted line, transcripts by a blue solid line, and proteins by an orange solid line. (a) Profiles of a xylosylated glucan epitope (GbXG-XX-2-C) and correlated representative transcript and protein of a xyloglucan xylosyltransferase (GbXXT2-1Dt). (b) Profiles of a galactosylated xyloglucan epitope (GbXG-XLX-1-C) and correlated representative transcript and protein of a xyloglucan galactosyltransferase (GbMUR3-2Dt). (c) Profiles of a xylan backbone epitope (GbXyl5-1-C) and correlated representative transcripts and proteins of a xylosyltransferase (GbIRX9-4At). (d) Profiles of a methylated-glucuronoxylan epitope (GbMeGlcA-2-K) and correlated representative transcripts and proteins of a xylosyltransferase (GbIRX10-1At) and a transferase involved in reducing end synthesis (GbFRA8-1At). (e) Profiles of a demethylated-homogalacturonan epitope (GbHG-Bbde-C) and correlated representative transcript and protein of a galacturonosyltransferase (GbGAUT4-2Dt). (f) Profiles of a methylated-homogalacturonan epitope (GbHG-BBMe-K) and correlated representative transcript and protein of a methyltransferase (GbGAUT8-1At). (g) Profiles of a β-1,4-galactan secondary backbone epitope of RG-I (GbGAL4-BB-C) and correlated representative transcript and protein of a β-1,4-galactan synthase (GbGALS1-2Dt). (h) Profiles of a branched-Rhamnogalacturonan-I epitope (GbRG-I-Gal3-7-K) and correlated representative transcript and protein of an arabinogalactan galactosyltransferase (GbAG:GALT8-3Dt). (Refer to Table S6 for the full list of correlated and non-correlated polysaccharides, transcripts and proteins).

The analysis revealed that many of the xylan-related glycosyltransferases from both the proteome and transcriptome data correlated with the profiles of xylan backbone (Xyl-BB) and branched xylan epitopes (GbXyl-GlcA and GbXyl-MeGlcA) (Figure 2c and 2d; Table S6). GbIRX9-3At xylosyltransferase was found significantly correlated to the profiles of Xyl-BB (Figure 2c; Table S6). Many of the glycosyltransferases include GbIRX9-3At/Dt, GbIRX10-1At/Dt, GbIRX15-2At/Dt, GbFRA8-1At/Dt, GbGUX1-1At/Dt, GbGUX2-2Dt, and GbGXMT3-1At/Dt significantly correlated with GbXyl-MeGlcA epitopes (Figure 2d; Table S6). Also, a few methyltransferases (GbRWA1-1At/Dt, GbRWA1-4At/Dt) and two acetyltransferases (GbTBL32-1Dt, GbTBL32-2Dt) highly correlated with GbXyl-MeGlcA epitopes (Table S6). Analysis showed that the GbHG epitopes profile correlated with the transcripts and proteins of GbGAUT4-2Dt, GbQUA2-2At, and GbTBR-5At/Dt (Figure 2e and 2f; Table S6). Similarly, the profile of GbGal4-BB epitopes matched with the GbGALS1-2Dt enzyme and GbRG-I-Gal3-7-K epitope matched with the GbAG:GALT8-3At/Dt enzymes (Figure 2g and 2h; Table S6).

#### Glycome profiling: heat maps, and self-organizing maps (SOM) of diverse epitope patterns of polysaccharides in *G. hirsutum* (*Gh*) and *G. barbadense* (*Gb*)

To characterize polysaccharide epitope compositions in *Gh* and *Gb* fibers, ELISA absorbance data from glycome profiling of *Gh* and *Gb* fiber at 6-25 DPA (Figure S1; Table S3) were visualized as heat maps and grouped into SOM clusters. ELISA data for each epitope originated from two different fractions of the same sample, namely the buffer-soluble and alkali-soluble fractions (Table S3). Accordingly, epitopes were labeled with a suffix: “-C” for buffer-soluble fractions (50mM CDTA:50mM ammonium oxalate buffer extracts) and “-K” for alkali-soluble fractions (4M KOH extracts). ELISA data from *Gh* and *Gb* were analyzed together, but the identities of all 71 epitopes were maintained separately for both species.

Heat maps were generated separately for buffer-soluble polysaccharide and alkali-soluble polysaccharide fractions from both species (Figure 3). In these heat maps, epitopes are arranged one-to-one between *Gh* and *Gb* to highlight the differential distribution of epitopes during fiber development (Figure 3). The heat maps of buffer-soluble polysaccharides showed that many of the highly branched Rhamnogalacturonan-I (RG-I), arabinogalactan (AG), xyloglucan (XG), and some xylan (Xyl) epitopes were present in lower amounts in *Gb* and also their amount dropped down at earlier stages of fiber development in *Gb* in comparison with *Gh* (Figure 3a). The heat maps of alkali-soluble fractions showed that many of the xyloglucan and xylan epitopes are present at high levels and in comparable amounts between both species. In addition, the heat maps clearly showed that the profile patterns and content of some of the xyloglucan (XG-XX-1-K, XG-XX-2-K, XG-XLX-1-K, XG-XLX-2-K, XG-XLX-3-K, XG-XnLG-K), xylan (Xyl-2Ar-1-K, Xyl-2Ar-2-K, Xyl-3Ar, Xyl-MeGlcA-2-K, Xyl-3-1-K), and of the RG-I (RG-I-1-4-K, RG-Ia-K) epitopes were prominently different between two species (Figure 3b).

**Figure 3.**
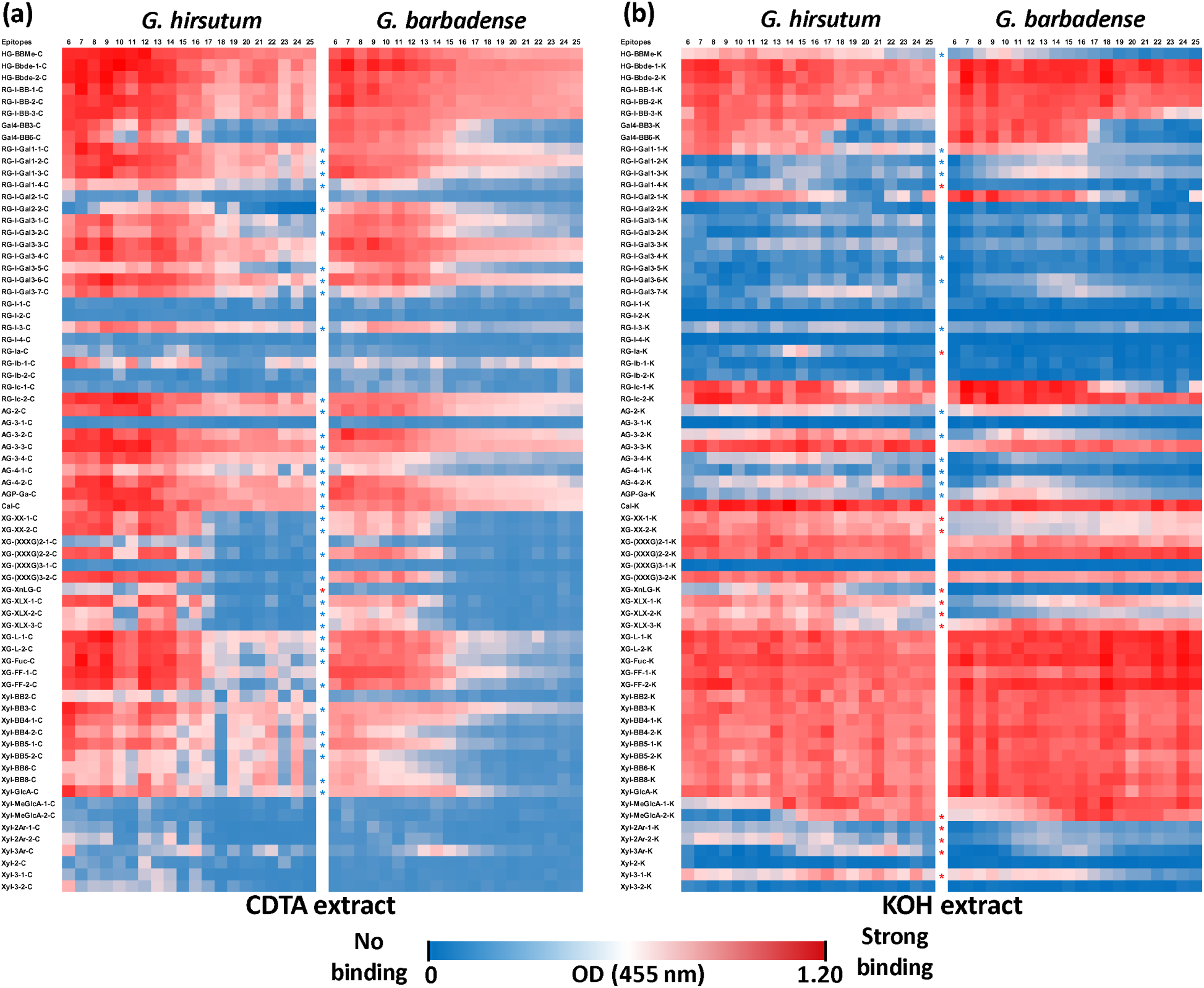
Heat maps of glycome profiled 71 different polysaccharides epitopes in fiber tissue of two cotton species during development (6 - 25 DPA). (a) Heat maps of glycome profiled epitopes from the buffer soluble (50mM CDTA-50mM Ammonium oxalate extract) polysaccharides fractions of G. hirsutum and G. barbadense. Data represent averaged optical density (OD) values from three biological replicates. (b) Heat maps of glycome profiled epitopes from the alkali soluble (4M KOH extract) polysaccharide fractions of G. hirsutum and G. barbadense. Data represent averaged OD from three biological replicates.

Epitopes from either buffer-soluble (50mM CDTA-50mM Ammonium oxalate extract) or alkali-soluble (4M KOH extract) fractions are denoted by the suffix “-C” and “-K”, respectively. Epitopes exhibiting similar patterns, but varying in content between the two species are marked with blue stars, while epitopes with different patterns between the species are marked with red stars. Detailed glycan epitope binding specificities of the 71 monoclonal antibodies used in this study are provided in Table S2.

Next, the glycome profiles from *Gh* and *Gb* were combined and clustered using an unbiased SOM approach to investigate the dynamics of epitope compositions across fiber development. Totally 12 SOM cluster groups were used to identify similarities and differences of polysaccharide epitope profiles between the two species (cluster G1-C to G12-C for buffer-soluble polysaccharide fractions and cluster G1-K to G12-K for alkali-soluble polysaccharide fractions) (Figure 4; Tables S7 and S8). The SOM profiles for buffer-soluble polysaccharides showed that most polysaccharide epitopes had higher contents at 6 DPA, and then gradually or rapidly reduced to a lower content at 25 DPA (Figure 4A). Epitopes in clusters G8-C, G11-C, and G12-C remained low throughout all DPAs. SOM clustering of alkali-soluble polysaccharides showed greater variability in epitope patterns, with some increasing or decreasing gradually or rapidly, while others peaked between 12 and 16 DPA during fiber development (Figure 4C). Members present in each SOM cluster are provided in Table S7.

**Figure 4.**
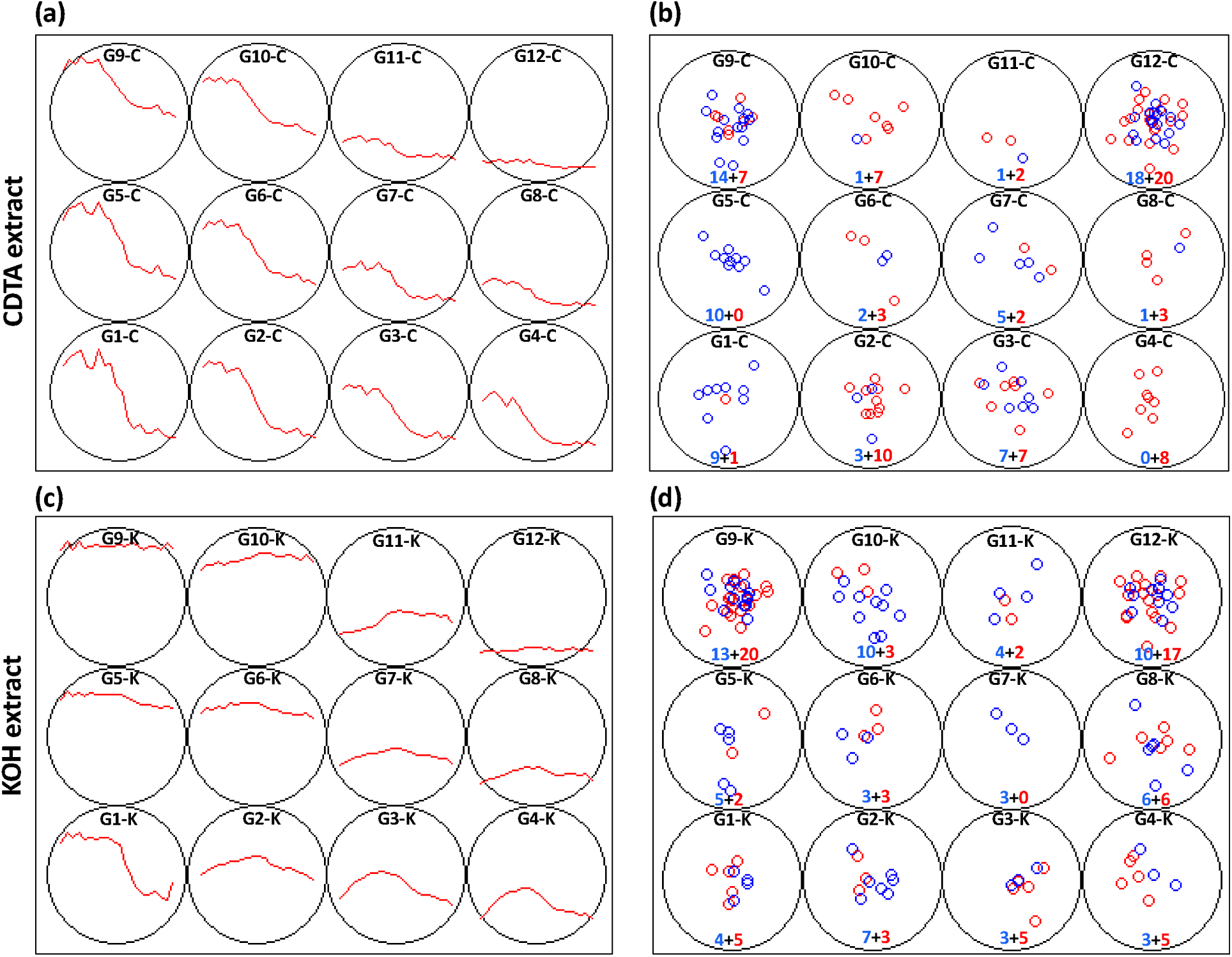
Self-organizing maps (SOM) of glycome profiled polysaccharide epitope patterns in cotton fibers from two species during development (6 - 25 DPA). (a) SOM groups (G1-C to G12-C) representing the glycome epitope profiles of buffer soluble polysaccharides (50mM CDTA-50mM Ammonium oxalate extract). (b) Distribution of epitopes from G. hirsutum (blue circles) and G. barbadense (red circles) in each SOM group shown in (a). (c) SOM groups (G1-K to G12-K) representing the glycome epitope profiles of alkali-soluble polysaccharide (4M KOH extract). (d) Distribution of epitopes from G. hirsutum (blue circles) and G. barbadense (red circles) in each SOM group shown in (c). Profiles in (a) and (c) represent averaged data from three biological replicates. In (b) and (d), each colored circle represents a specific epitope, and the numbers indicate the number of epitopes assigned to each group.

Both the SOM and heat maps provided complimentary details of the epitope patterns and their differential distribution between the fibers of the two species during development (6 - 25 DPA). These data allowed us to group epitopes into three major “categories” (Figures 5 – 7; Table S8), namely 1) polysaccharide epitopes that are similar in content and no temporal variability between the two species, 2) polysaccharide epitopes that have similar profiles through all 20 DPAs, but vary in their content between two species, and 3) polysaccharide epitopes that have more noticeable differences in their content and profiles between two species.

**Figure 5.**
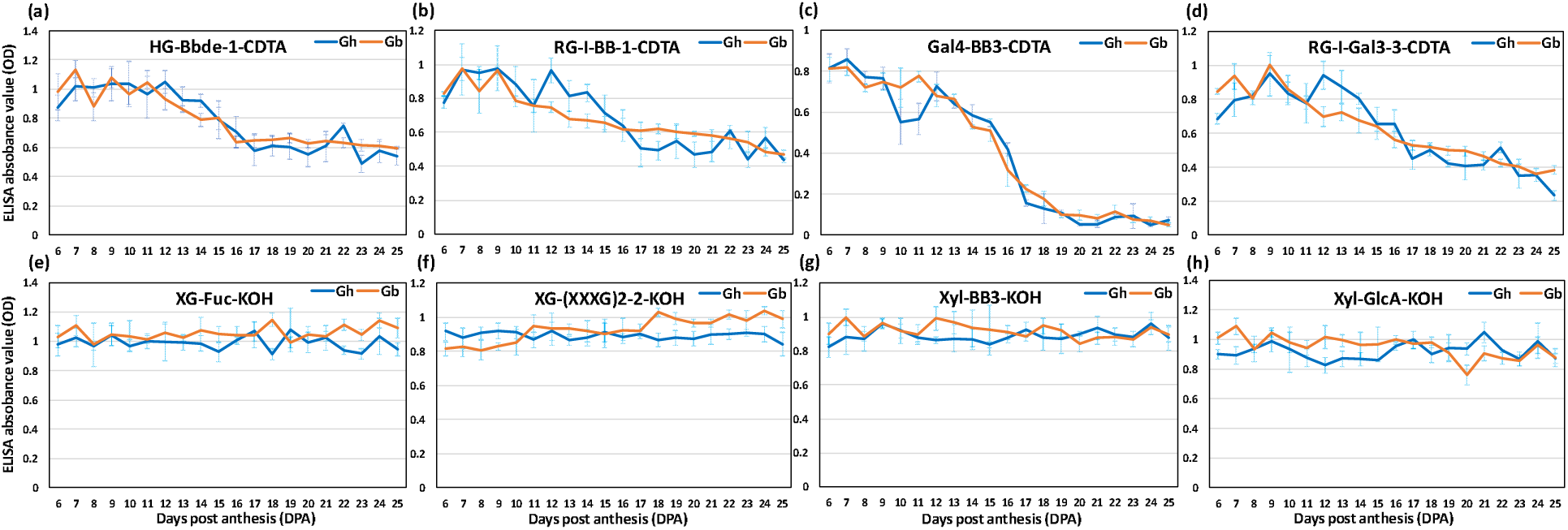
Glycome profiles of polysaccharide epitopes with equal content and same pattern between G. hirsutum (Gh) and G. barbadense (Gb) fiber across development (6 - 25 DPA). (a - h) Representative profiles of polysaccharide epitopes listed in Table S8A are shown here. The profiles of Gh (blue) and Gb (orange) show averaged optical density (OD) values of three biological replicates with standard error bars. Epitopes from either buffer-soluble (50mM CDTA-50mM Ammonium oxalate extract) or alkali-soluble (4M KOH extract) polysaccharide fractions are denoted by the suffix “-CDTA” and “-KOH”, respectively.

The first categorical group includes 44 polysaccharide epitopes with comparable contents and similar profiles in the two species, indicating no significant difference between some of the polysaccharide epitopes in their fibers (Figure 5; Table S8A). This group contains epitopes present in HGs (both methyl esterified and de-esterified), RG-I-BB (RG-I backbone epitopes), Gal4-BB (secondary galactan backbone of RG-I), highly branched RG-I (RG-I-Gal2, RG-I-Gal3s), xyloglucans (XG-(XXXG)2, XG-(XXXG)3, XG-L, XG-FF), and xylans (Xyl-BBs, Xyl-GlcA, Xyl-MeGlcA-1). Most of the epitopes in this category, from *Gh* or *Gb* were found in the same SOM groups. For instance, buffer-soluble HG epitopes (HG-Bbde-1-C/-2-C, HG-BBMe-C) and RG-I epitopes (RG-I-BB-1-C/-2-C/-3-C, RG-I-Gal3-3-C) of both *Gh* and *Gb* were present in the same SOM group G9-C (Figure 5; Table S8A). Similarly, some alkali-soluble HG epitopes (HG-Bbde-1-K, HG-Bbde-2-K), RG-I epitopes (RG-I-BB-1-K, RG-I-Gal2-1-K, RG-Ic-2-K), xyloglucan epitopes (XG-(XXXG)2-2-K, XG-FF-2-K, XG-Fuc-K, XG-L-1-K, XG-L-2-K), and xylan epitopes (Xyl-BB3-K, Xyl-GlcA-K) from both species were assigned in the same G9-K SOM cluster. Many other polysaccharide epitopes from both species were also grouped in the same SOM groups, e.g., RG-I-BB-3-K, Gal4-BB3-C, Gal4-BB3-K, Gal4-BB6-K, RG-Ic-1-C, RG-I-Gal2-1-K, RG-I-Gal3-5-C, RG-I-Gal3-7-K, Xyl-MeGlcA-1-K, and Cal-K. SOM analysis revealed subtle differences in overall patterns and abundance of epitopes between the species. For example, many of the xylan backbone epitopes (Xyl-BBs) from *Gh* were found in G10-K SOM group, whereas the same epitopes from *Gb* were found in the neighboring G9-K SOM group, although profiles in these two SOM groups were not statistically different (Figure 5; Table S8A).

The other 48 epitopes fell into the second categorical group, in which they had the same profiles, but their contents significantly varied between *Gh* and *Gb* (Figure 6; Table S8B). Many of the highly branched RG-I, arabinogalactan (AGs), xyloglucan, and some xylan epitopes were included in this group. Despite having similar profile, these epitopes showed significant content differences between the two species, and the SOM analysis effectively distinguished them based on their quantitative differences. For example, in *Gh*, six xyloglucan epitopes (XG-XX-1-C, XG-XX-2-C, XG-(XXXG)2-2-C, XG-(XXXG)3-2-C, XG-FF-2-C, XG-XLX-1-C) and three highly branched RG-I epitopes (RG-I-Gal3-1-C, RG-I-Gal3-2-C, RG-I-Gal3-7-C) were assigned to the G1-C group (Figure 6; Table S8B). In contrast, the same above epitopes from *Gb* were distributed to the G2-C and G4-C groups. Similarly, the G5-C group contained only polysaccharide epitopes from *Gh*, such as branched RG-I and xyloglucan epitopes, whereas the same epitopes from *Gb* were predominantly distributed to the G2-C and G10-C groups. Interestingly, only four epitopes out of 48 were present in higher amounts in *Gb* in comparison with *Gh*, whereas the remaining 44 epitopes showed higher presence in *Gh* relative to *Gb* (Table S8B). Overall, glycome profiling showed that *Gb* has significantly lower amounts of highly branched RG-I, arabinogalactan, and xyloglucan epitopes in comparison with *Gh* (Figure 5; Table S8B).

**Figure 6.**
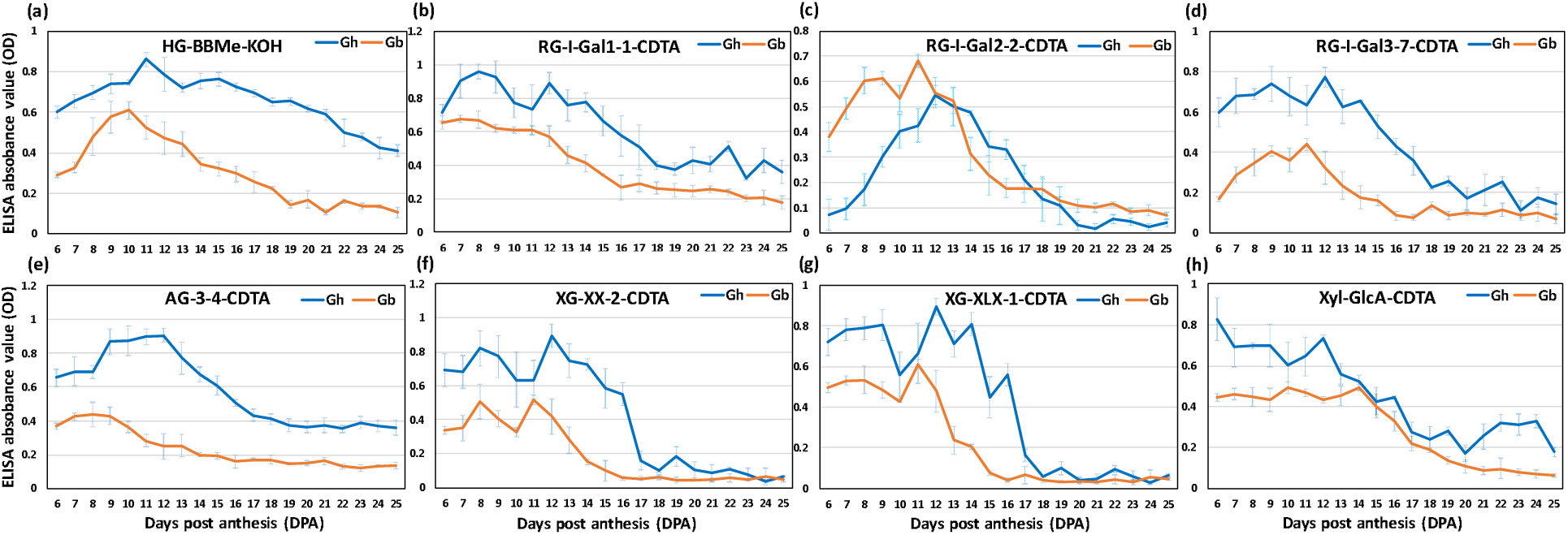
Glycome profiles of polysaccharide epitopes with the same pattern, but varying content between G. hirsutum (Gh) and G. barbadense (Gb) fiber across development (6 - 25 DPA). (a - h) Representative profiles of polysaccharide epitopes listed in Table S8B are shown here. The profiles of Gh (blue) and Gb (orange) show averaged optical density (OD) values of three biological replicates with standard error bars. Epitopes from either buffer-soluble (50mM CDTA-50mM Ammonium oxalate extract) or alkali-soluble (4M KOH extract) polysaccharides fractions are denoted by the suffix “-CDTA” and “-KOH”, respectively.

Comparative heat maps and SOM analysis clearly differentiated 14 polysaccharide epitopes with different profile patterns among the two species, forming the third categorical group (Figure 7; Table S8C). Interestingly, three epitopes, RG-Ia-K (present in the group G4-K for *Gh*, G12-K for *Gb*), XG-XnLG-K (present in G3-K for Gh, G12-K for Gb), and RG-I-Gal1-4-K (present in G4-K for *Gh*, G11-K for *Gb*), where their contents were found low and profiles were flat in *Gb* through 6 to 25 DPA. On the contrary, in *Gh*, the profiles of these polysaccharide epitopes showed a semi-bell-shaped curve and peaked in the middle of the fiber development. The content of five xyloglucan epitopes, XG-XX-1 (present in G6-K for *Gh*, G2-K for *Gb*), XG-XX-2 (present in G5-K for *Gh*, G2-K for *Gb*), XG-XLX-1 (present in G6-K for *Gh*, G2-K for *Gb*), XG-XLX-2 (in G2-K for *Gh*, G11-K for *Gb*), and XG-XLX-3 (in G1-K for *Gh*, G6-K were *Gb*), and two xylan epitopes, Xyl-2Ar-1 (in G4-K for *Gh*, G8-K for *Gb*), and Xyl-2Ar-2 (in G3-K for *Gh*, G8-K for *Gb*) were lower in content in *Gb* at the earlier DPAs but increased gradually to match the amount of *Gh* at later stages (Figure 7; Table S8C). A reverse profile was seen for a xylan epitope, Xyl-MeGlcA-2 (present in G2-K for *Gh*, G10-K for *Gb*), for which the content was lower in *Gh* and increased gradually to equal the amount present in *Gb* at the later DPAs. In *Gb*, the amount of two of the xylan epitopes, Xyl-3Ar (present in G11-K for *Gh*, G8-K for *Gb*) and Xyl-3-1 (in G2-K for *Gh*, G3-K for *Gb*) was comparable to *Gh* at earlier stages but drastically reduced after 15 DPA to reach a significantly lower level in *Gb* by 25 DPA.

**Figure 7.**
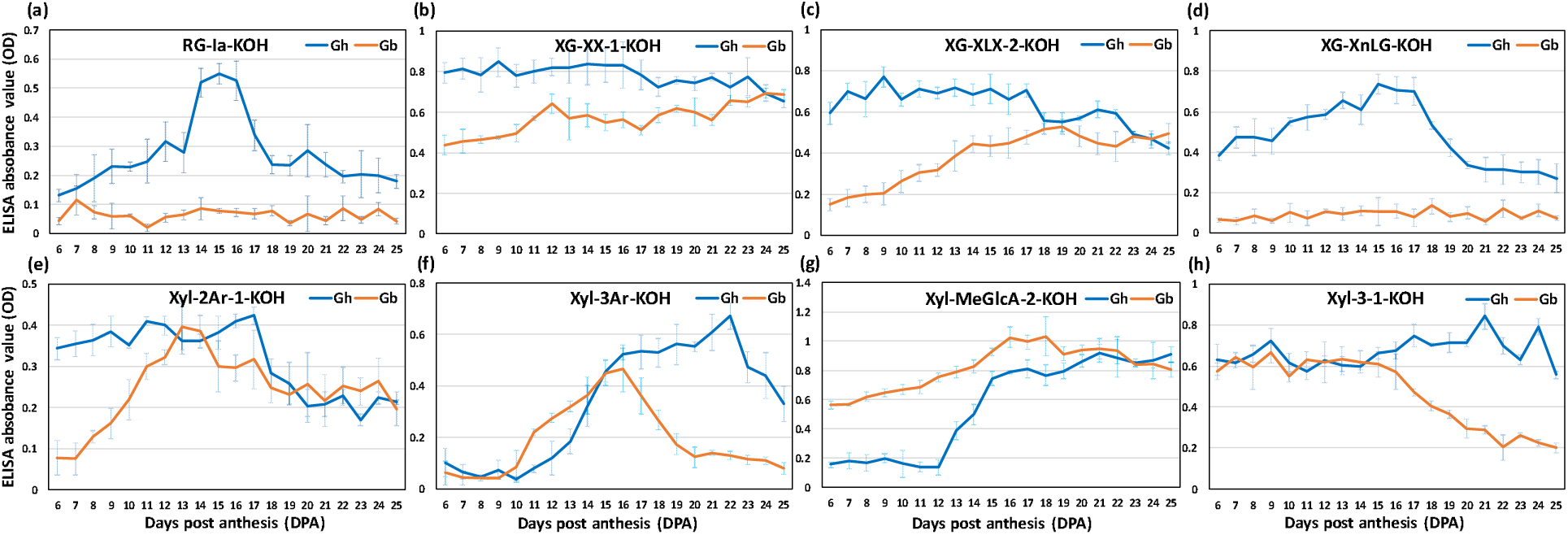
Glycome profiles of polysaccharide epitopes with different pattern between G. hirsutum (Gh) and G. barbadense (Gb) fiber across development (6 - 25 DPA). (a - h) Representative profiles of polysaccharide epitopes listed in Table S8C are shown here. The profiles of Gh (blue) and Gb (orange) show averaged optical density (OD) values of three biological replicates with standard error bars. Epitopes from alkali soluble (4M KOH extract) fractions are denoted by the suffix “-KOH”.

There are about 34 epitopes detected at very low levels and showed noisy profiles, and therefore were not analyzed further; these epitopes were mainly present in SOM groups, G12-C and G12-K (Figure 4; Table S8D).

#### Differentially expressed transcripts of glycosyltransferases between *G. hirsutum* (*Gh*) and

##### G. barbadense (Gb)

Based on the gene sequences of known *Arabidopsis* polysaccharide-synthesizing enzymes, transcript data of the corresponding orthologous cotton genes were extracted from the whole transcriptomic data of both cotton species (Table S4; Swaminathan et al., 2024). Subsequently, the comparisons of transcript expression levels of different enzymes between *Gh* and *Gb* and their sub-genomes (At and Dt) were estimated.

##### Differentially expressed transcripts of cellulose synthases (CESAs)

Two types of cellulose synthases (CESAs), the PCW CESAs (CESAs 1/3/6) and SCW CESAs (CESAs 4/7/8), are known to be involved in synthesizing cellulose microfibrils of PCW and SCW, respectively (Kim et al., 2019). Using *Arabidopsis* gene homology analysis, four CESA1, six CESA3, and 14 CESA6 and two each for CESA4, CESA7 and CESA8, including from both the A sub-genome (At) and D sub-genome (Dt) were identified in our cotton transcripts (Table S4). Further, we compared the transcripts of these CESAs on a one-to-one basis and found that two of the PCW CESAs (*CESA3-C-Dt* and *CESA6-D-At*) and ten of the SCW CESAs (*CESA4-A-At* & *-Dt*, *CESA4-B-At* & *-Dt*, *CESA7-A-At* & *-Dt*, *CESA7-B-At* & *-Dt*, and *CESA8-B-At* & *-Dt*) (including both A and D genomes) were differentially expressed between *Gh* and *Gb* (Figure 8; Table S4). The PCW *CESAs* transcripts from *Gb* (*CESA3-C-Dt* and *CESA6-D-At*) were expressed higher in *Gb* compared to *Gh*, whereas levels of all other PCW *CESAs* transcripts were similar in both species. In contrast, the ten differentially expressed SCW *CESAs* showed higher transcript levels in *Gh* than in *Gb*. Interestingly, there was a two to three-day delay in the expression of all ten SCW CESAs in *Gb* fiber in comparison with *Gh*. In *Gh*, the transcript level of SCW *CESAs* started to rapidly increase around 13 DPA, whereas in *Gb* such rapid increase began around 15 DPA.

**Figure 8.**
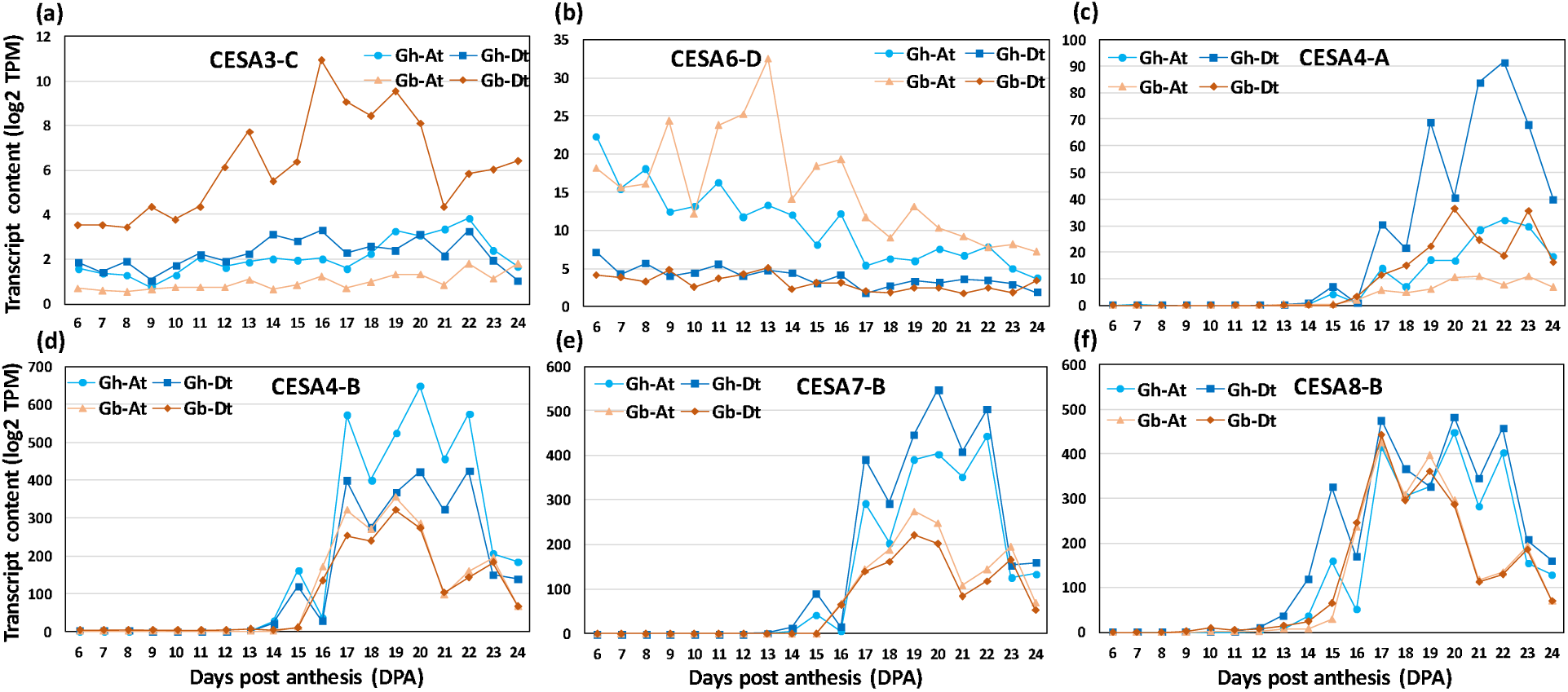
The profiles of cellulose synthase (CESAs) transcripts differentially expressed in G. hirsutum (Gh) and G. barbadense (Gb) fiber during development (6 - 24 DPA). (a, b) Transcript abundance profiles of primary CW (PCW) synthesizing CESAs. (c – f) Transcript abundance profiles of secondary CW (SCW) synthesizing CESAs. Log2-transformed transcript per million (TPM) values of A sub-genome (At) and D sub-genome (Dt) of both the species are shown in the plot.

##### Differentially expressed transcripts of xyloglucan-synthesizing glycosyltransferases

Next, we analyzed glycosyltransferases known to be involved in xyloglucan biosynthesis. Well-characterized xyloglucan-synthesizing enzymes from *Arabidopsis* are xyloglucan β-1,4-glucan synthases (cellulose synthase-like-C enzymes, CSLCs; CSLC4/5/6/8/12), xyloglucan α-1,6-xylosyltransferases (XXT1/2/3/4/5), xyloglucan β-1,2-galactosyltransferases (MUR3 and XLT2), and a α-1,2-L-fucosyltransferase (FUT1) (Figure S1) (Julian & Zabotina, 2022). These CSCLs involved in the synthesis of the glucan backbone; XXTs add xylose residues to the specific glucoses, MUR3/XLT2 galactosylates specific xyloses, and FUT1 fucosylates the galactose residues. By homology search, we found that cotton has 16 *CSLCs*, 10 *XXTs*, 8 *MUR3s*, 4 *XLT2s*, and 4 *FUT1s* from both the A and D sub-genomes (Table S4; Swaminathan et al., 2024). The comparative analysis of *Gh* and *Gb* from both the A and D genomes showed that for all differentially expressed xyloglucan-synthesizing enzymes, *Gh* has a higher transcript level than *Gb* (Figure 9; Table S4). The differentially and highly expressed glycosyltransferases in *Gh* were, five CSLCs (*CSLC04-1At* & *-1Dt, CSLC05-2At & -2Dt, CSLC12-2Dt*, four XXTs (*XXT2-2At* & *-2Dt, XXT3-1At* & *-1Dt*), three MUR3s (*MUR3-1At, MUR3-4At* & *-4Dt*), and two XLTs (*XLT-1At* & *-1Dt*).

**Figure 9.**
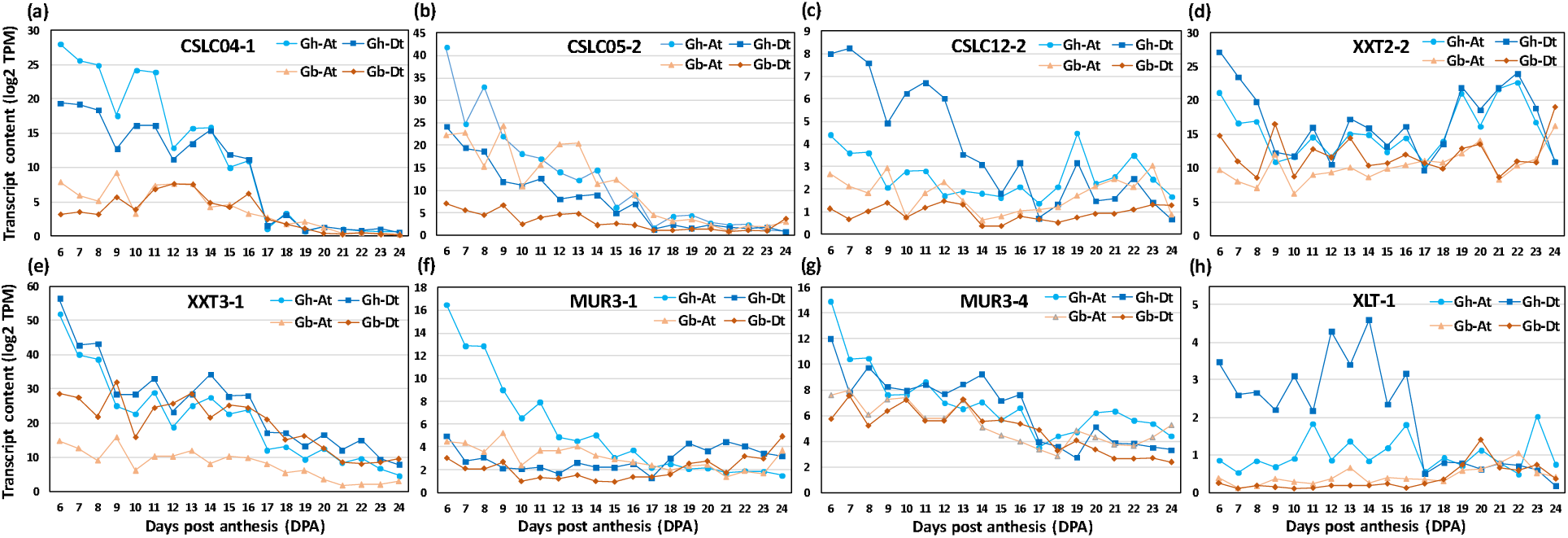
The profiles of xyloglucan-synthesizing glycosyltransferase transcripts differentially expressed in G. hirsutum (Gh) and G. barbadense (Gb) fiber during development (6 - 24 DPA). (a – h) Transcript abundance profiles of representative cellulose synthase-like-C enzymes (CSLCs); UDP-Xyl: xyloglucan α-1,6-xylosyltransferases (XXTs), and xyloglucan ß-1,2-galactosyltransferases (MUR3 and XLT2). Log2-transformed transcript per million (TPM) values of A sub-genome (At) and D sub-genome (Dt) of both the species are shown in the plot.

##### Differentially expressed transcripts of xylan-synthesizing glycosyltransferases

Xylan-synthesizing glycosyltransferases identified so far are xylan β-1,4-xylosyltransferases (IRXs; irregular xylem), xylan α-glucuronyltransferases (GUXs), glucuronoxylan methyltransferase-like proteins (GXMT), and xylan arabinosyltransferases (XATs) (Figure S1) (Smith et al., 2017; Chen et al., 2020a). IRXs are responsible for xylan backbone synthesis. GUXs are involved in adding α-1,2-d-glucuronic acid (GlcA) to the xylan backbone, and GXMTs methylate glucuronic acids in the side chains of xylans. XATs are involved in adding arabinose residues to xylan backbones, while O-acetyltransferases (ESK/RWA/TBL) acetylate the backbone. In addition, there are other enzymes proposed to synthesize the specific reducing end oligosaccharide of the xylans (FRA8-1/PARVUS/IRX8), and the formation of which leads to the termination of xylan backbone elongation. Homology analysis using known *Arabidopsis* genes revealed 30 *IRXs*, 16 *GUXs*, 22 *GXMTs*, 40 O-acetyltransferases (*ESK/RWA/TBL*) and 14 GTs predicted to synthesize the reducing end of xylan (*FRA8-1/PARVUS/IRX8*), from both A and D cotton genomes (Table S4).

Comparative transcript analysis of xylan backbone-synthesizing glycosyltransferases demonstrated that 17 out of 30 IRXs (*IRX9-3Dt, IRX10-1At* & *-1Dt, IRX10-2At* & *-2Dt, IRX10-3At* & *-3Dt, IRX14-1At* & *-1Dt, IRX14-2At* & *-2Dt, IRX15-1At* & *-1Dt, IRX15-2Dt, IRX15-3At* & *-3Dt, IRX15-4Dt*) were present in higher amount in *Gh* in comparison with *Gb* (Figure 10; Table S4). Interestingly, only two IRXs (*IRX9-2At* & *-2Dt*) were found to be highly expressed in *Gb*. Ten O-acetyltransferases (*ESK1-1At* & *-1Dt, RWA1-1Dt, RWA1-3At, RWA1-4At* & *-4Dt, TBL3-1At* & *-1Dt, TBL3-2At* & *-2Dt*) were more highly expressed in *Gh* than in *Gb*, while only one was more highly expressed in *Gb* (*TBL30-3Dt*) than in *Gh*. Among the enzymes proposed to synthesize the reducing end oligosaccharide in xylan, ten glycosyltransferases (*FRA8-1Dt, PARVUS-1At* & *-1Dt, PARVUS-2At* & *-2Dt, PARVUS-3At, PARVUS-4Dt, IRX8-1At, IRX8-2At* & *-2Dt*) showed higher transcript levels in *Gh* relative to *Gb*. However, none of these transcripts were found to be higher in *Gb* than in *Gh*. Among the enzymes involved in branching of the xylan backbone, two GUXs (*GUX1-2At, GUX2-1At*) transcripts were highly expressed in *Gh*, whereas four were more highly expressed in *Gb* (*GUX1-1At, GUX2-2At* & *-2Dt, GUX3-1Dt*). Among GXMTs, four transcripts were more highly expressed in *Gh* (*GXMT4-3At* & *-3Dt, GXMT4-4At* & *-4Dt*), whereas three had higher levels in *Gb* (*GXMT2-1At, GXMT5-2At* & *-2Dt*). Among XATs transcripts, two were more highly expressed in *Gh* (*XAT-1At, XAT-3Dt*) and only one was more highly expressed in *Gb* than in *Gh* (*XAT-2At*).

**Figure 10.**
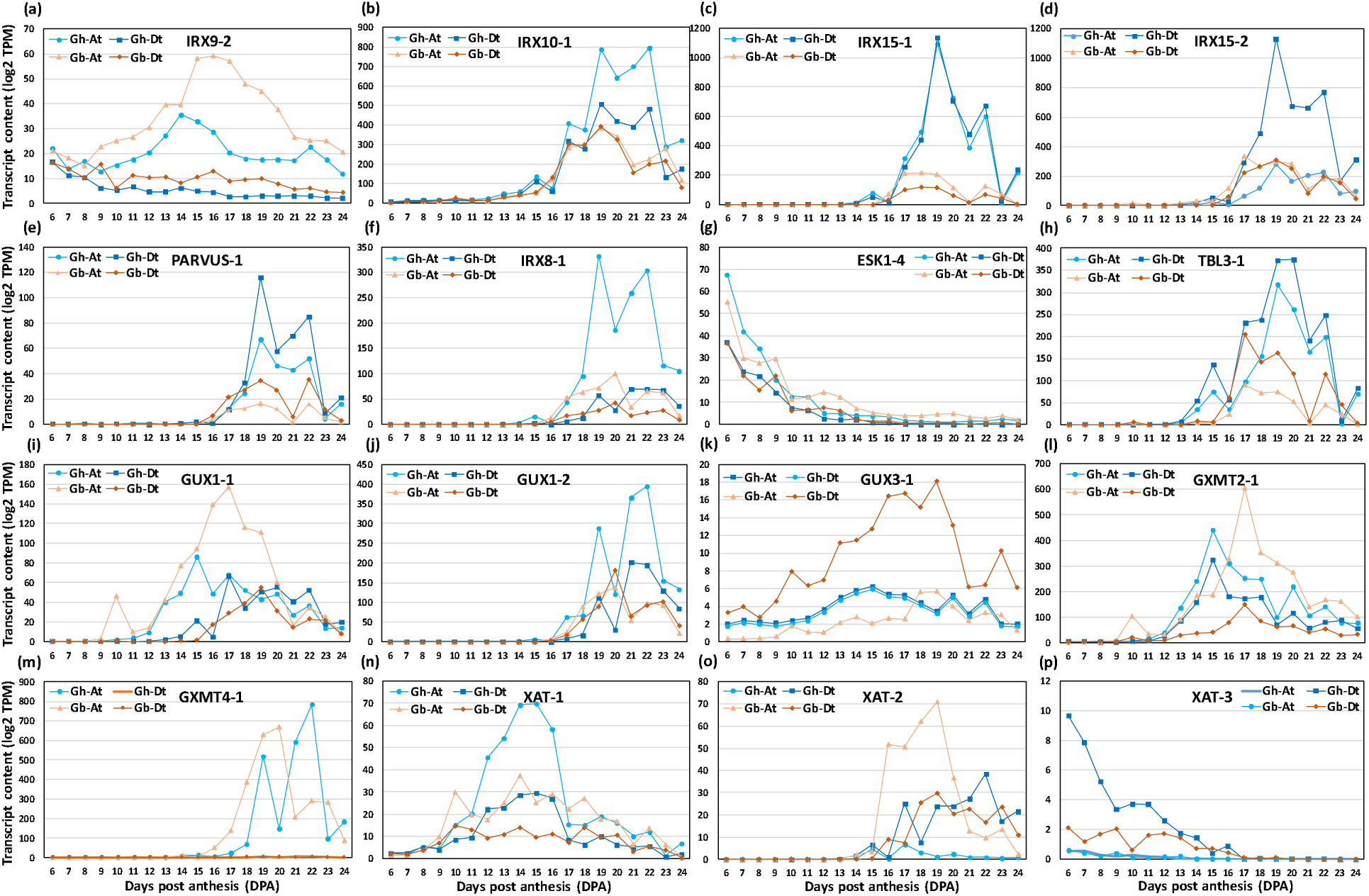
The profiles of xylan-synthesizing glycosyltransferase transcripts differentially expressed in G. hirsutum (Gh) and G. barbadense (Gb) fiber during development (6 - 24 DPA). (a – p) Transcript abundance profiles of representative xylan ß-1,4-xylosyltransferases (IRXs; irregular xylem), O-acetyltransferases (ESK/TBL), reducing end synthesizing glycosyltransferase (PARVUS), UDP-GlcA: xylan α-glucuronyltransferases (GUXs), glucuronoxylan methyltransferase-like proteins (GXMT), and xylan arabinosyltransferases (XATs). Log2-transformed transcript per million (TPM) values of A sub-genome (At) and D sub-genome (Dt) of both the species are shown in the plot.

##### Differentially expressed transcripts of homogalacturonan synthesizing glycosyltransferases

Synthesis of homogalacturonan (HG) is carried out by galacturonosyltransferase (GAUTs) and galacturonosyltransferase-like (GATLs) enzymes, which add galacturonic acid (GalA) residues to the growing chain of HG backbones. The methyl and acetyl groups are added to the HG backbone by methyl transferases (CGRs/QUAs) and O-acetyltransferases (TBLs/TBRs), respectively (Atmodjo et al., 2013; Engle et al., 2022) (Figure S1).

Homology search of *Arabidopsis* HG-related GTs revealed that there are 48 *GAUTs*, 24 *GALTs*, 24 methyltransferases (*CGRs/QUAs*), and 14 acetyltransferases (*TBLs/TBRs*) genes expressed in cotton fibers, which includes both A and D sub-genomes (Table S4; Swaminathan et al., 2024). Comparative transcript analysis showed that ten GAUTs were expressed at a higher level in *Gh* than in *Gb* (*GAUT1-4Dt, GAUT3-1At, GAUT4-1Dt, GAUT6-5At, GAUT8-2Dt, GAUT9-1At, GAUT12-1At* & *-1Dt, GAUT12-2At* & *-2Dt*), whereas four were more highly expressed in Gb (*GAUT7-1Dt, GAUT8-1Dt, GAUT15-1At* & *-1Dt*) (Figure 11; Table S4). Among GATLs, ten were found to be highly expressed in *Gh* (*GATL1-3At* & *-3Dt, GATL1-4At* & *-4Dt, GATL1-5At* & *-5Dt, GATL2-2Dt, GATL3-1Dt, GATL7-2Dt, GATL10-1Dt*), whereas only one was in *Gb* (*GATL2-3Dt*). Among methyl transferases, six were highly expressed in *Gh* (*CGR2-2At* & *-2Dt, CGR2-4At, QUA2-1Dt, QUA2-2At, QUA3-1At*) and none were in *Gb*. Among acetyltransferases, the transcript levels of five were higher in *Gh* (*TBL3-1At* & *-1Dt, TBL3-2At* & *-2Dt, TBR-5At*) and only one was higher in *Gb* (*TBR-1Dt*) (Figure 11; Table S4).

**Figure 11.**
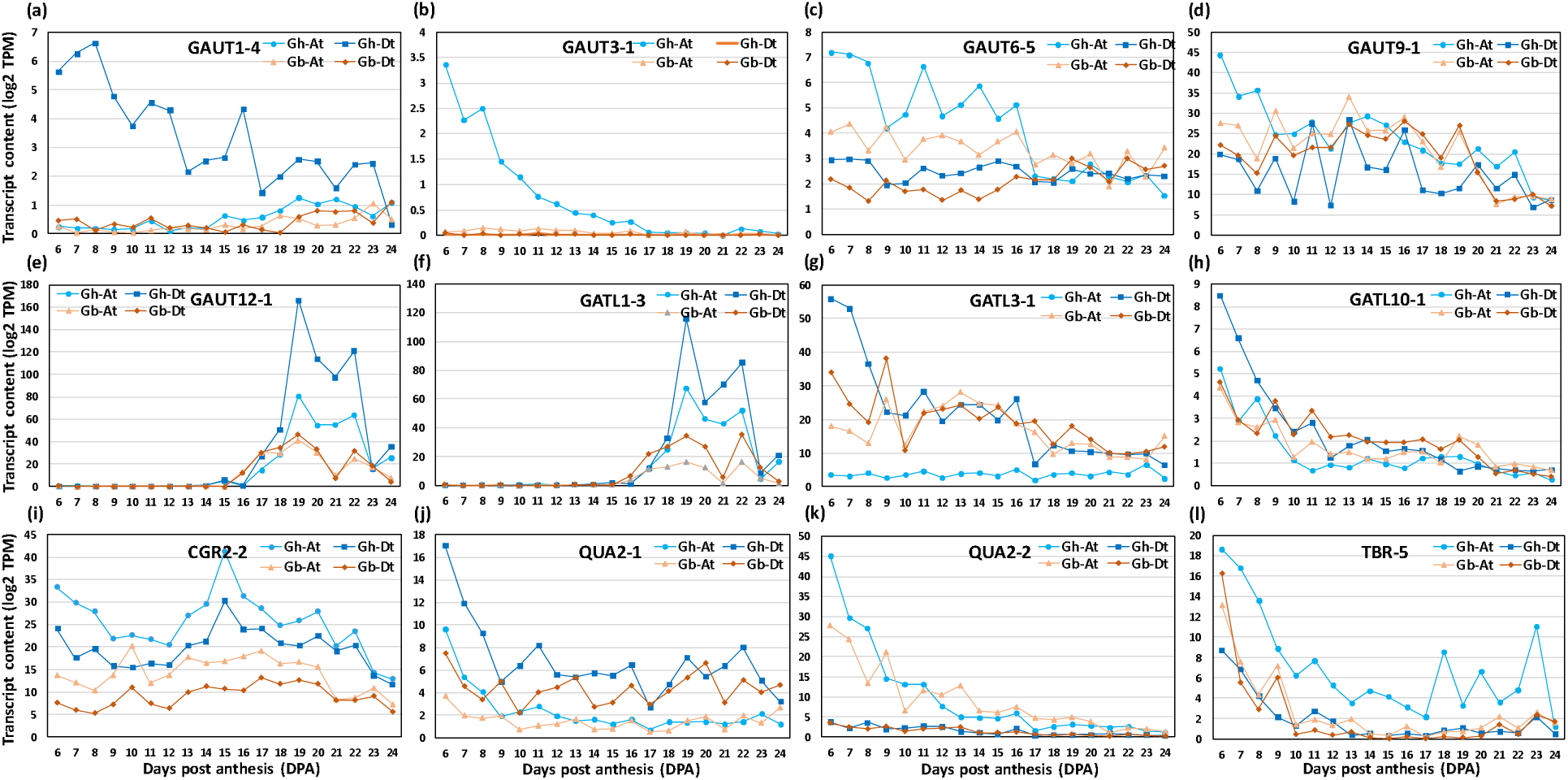
The profiles of homogalacturonan-synthesizing glycosyltransferase transcripts differentially expressed in G. hirsutum (Gh) and G. barbadense (Gb) fiber during development (6 - 24 DPA). (a – l) Transcript profiles of representative galacturonosyltransferases (GAUTs), galacturonosyl transferase-like enzymes (GATLs), methyltransferases (CGRs/QUAs), and acetyltransferases (TBRs). Log2-transformed transcript per million (TPM) values of A sub-genome (At) and D sub-genome (Dt) of both the species are shown in the plot.

##### Differentially expressed transcripts of Rhamnogalacturonan-I synthesizing glycosyltransferases

Rhamnogalacturonan-I (RG-I) is a highly branched pectin that has a complex structure. RG-I primary backbone is branched with the β-1,3-galactans, β-1,4-galactans, and arabinans, which constitute the secondary backbone of RG-I molecules. Further, the secondary backbone galactans are decorated with β-1,6-linked galactans, and arabinan side chains (Figure S1) (Atmodjo et al., 2013; Amos et al., 2022). RG-I:Galacturonosyltransferases (RG-I:GalATs) and RG-I:rhamnosyltransferases (RRTs) are known to synthesize the primary backbone of RG-I in *Arabidopsis*. The AG-GALTs and β-1,4-galactan synthases (GALSs) synthesize β-1,3-galactan and β-1,4-galactan secondary backbone of RG-I, respectively. The β-1,6-galactosyltransferases (GALT29A/GALT31A) and β-1,3-glucuronosyl transferases (GlcAT14) are involved in decorating secondary backbones of RG-I with β-1,6-linked galactans, and β-1,3-linked glucuronic acid, respectively (Showalter & Basu, 2016). The arabinan α-1,5-L-arabinosyltransferases (ARAD) add the arabinan backbone chain to the RG-I backbone. Homology searching of *Arabidopsis* genes involved in RG-I biosynthesis showed that there are 8 *RG-I:GalATs*, 34 *RRTs*, 6 *GALSs*, 34 *AG:GALTs*, 10 *GALT29A/GALT31As*, and 4 *ARAD*s in cotton fibers, from both A and D homeologs (Table S4).

Comparative transcriptome analysis of RG-I synthesizing glycosyltransferases revealed that three RG-I:GalATs were more highly expressed in *Gh* (*RGI:GalAT1-1Dt, RGI:GalAT1-3At* & *-3Dt* ) than in *Gb* and none in *Gb* in comparison with *Gh* (Figure 12; Table S4). Among RRTs, three were more expressed in *Gh* (*RRT1-2At, RRT4-1Dt, RRT9-2At*) and two were in *Gb* (*RRT1-4Dt, RRT9-1At*). Out of all GALS, the expression of one *GALS2-1Dt* was higher in *Gh*, while another *GALS1-2Dt* was more highly expressed in *Gb*. Among AG:GALTs, the transcript levels of five were higher in *Gh* (*AG:GALT1-1At, AG:GALT4-3At, AG:GALT5-2At, AG:GALT5-3Dt, AG:GALT8-2At*) and only *AG:GALT1-2Dt* had a higher expression level in *Gb*. Among *GALT29As/GALT31As*, five transcripts, *GALT29A-1At, GALT29A-2At, GALT31A-1At* & *-1Dt, GALT31A-2Dt* were more highly expressed in *Gh* in comparison with *Gb* and none in *Gb*. Among all GlcAT14s, eight were more highly expressed in *Gh* (*GlcAT14A-1At* & *-1Dt, GlcAT14A-3At* & *-3Dt, GlcAT14A-4At, GlcAT14A-5At* & *-5Dt, GlcAT14C-1Dt*) and none was present at a higher level in *Gb*. Among ARADs, two were more highly expressed in *Gh* (*ARAD1-1At* & *-1Dt*) and only *ARAD1-2Dt* was higher in *Gb* than in *Gh* (Figure 12; Table S4).

**Figure 12.**
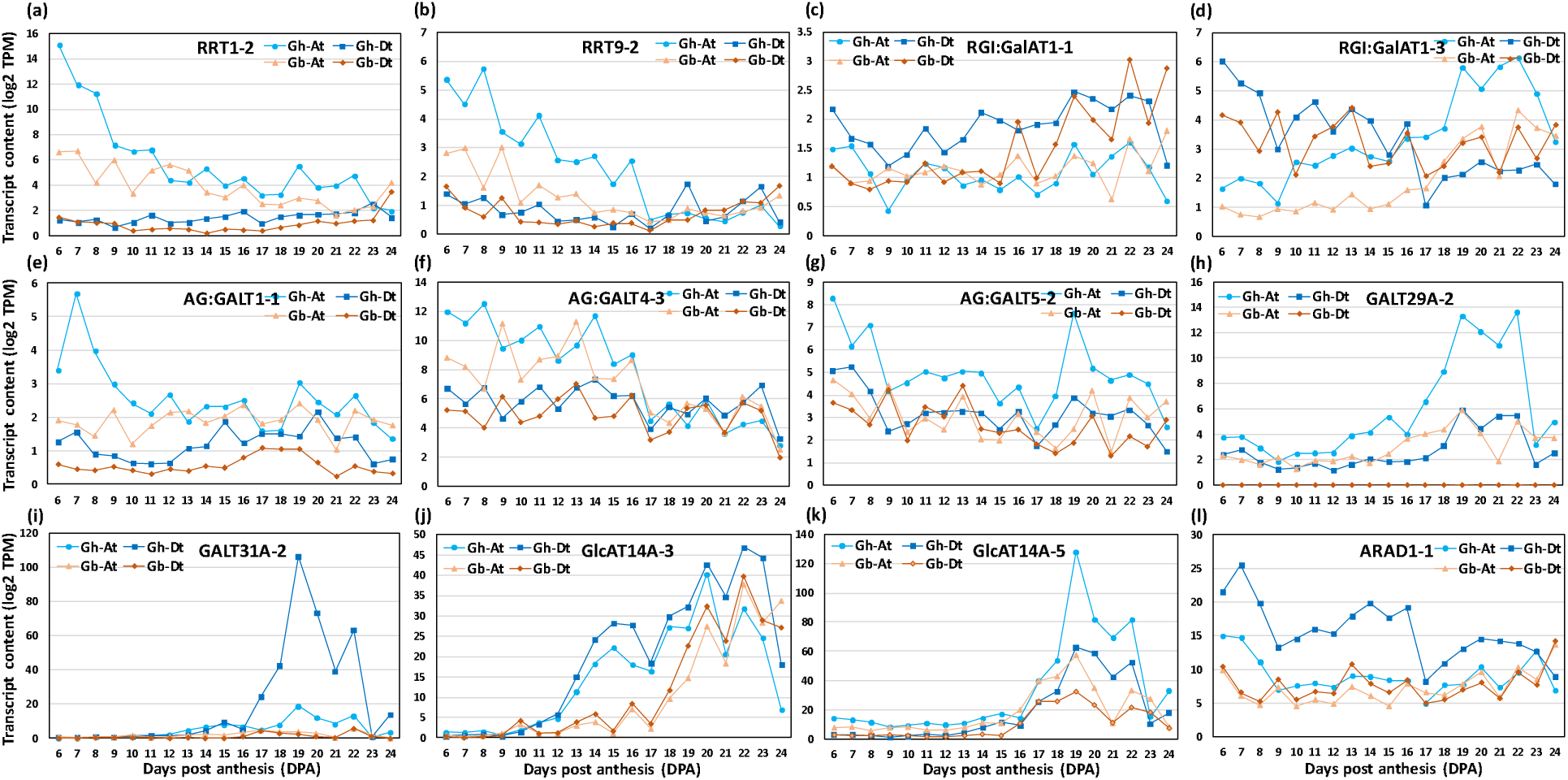
The profiles of RG-I-synthesizing glycosyltransferase transcripts differentially expressed in G. hirsutum (Gh) and G. barbadense (Gb) fiber during development (6 - 24 DPA). (a – l) Transcript profiles of representative RG-I:rhamnosyltransferases (RRTs), galacturonosyl transferases (RG-I:GalATs), ß-1,6-galactosyltransferases (GALT29A/GALT31A/AG:GALTs), ß-1,6-glucuronosyl transferases (GlcAT), and arabinan a-1,5-L-arabinosyltransferase (ARAD). Log2-transformed transcript per million (TPM) values of A sub-genome (At) and D sub-genome (Dt) of both the species are shown in the plot.

##### Differentially expressed transcripts of expansins

Recent studies highlighted the importance of CW-loosening protein expansin as one of the major players in fiber elongation (Sampedro and Cosgrove, 2005; Xu et al., 2013; Li et al., 2016; Lv et al. 2020). We compared the transcript levels of expansins from both *Gh* and *Gb* fibers. Levels of many of the expansion transcripts were higher in *Gb* than in *Gh* (Figure 13; Table S9). Thus, 14 expansins (*EXPA4-1Dt, EXPA4-2At* & *-2Dt, EXPA4-3At, EXPA4-5At* & *-5Dt, EXPA4-6At* & *-6Dt, EXPA8-2Dt, EXPA13-1At, EXPLB3-1At, EXPLA1-2At, EXP11-1At* & *-1Dt*) were highly expressed in *Gb*, whereas eight (*EXPA4-4At* & *-4Dt, EXPA8-2At, EXPA8-3Dt, EXPA15-1At, EXPA15-2Dt, EXPLB3-1Dt, EXPLA1-2Dt*) had higher transcript levels in *Gh* relative to *Gb*. In *Gb*, the transcript content of eight of the expansins (*EXPA4-5At* & *-5Dt, EXPA4-6At* & *-6Dt, EXPLB3-1At, EXPLA1-2At, EXP11-1At* & *-1Dt*) rapidly increased after 12 DPA and were present at higher levels at later DPAs. In the case of *Gh*, only two transcripts (*EXPLA1-2At* & *-2Dt*) were found to have similar expression patterns, but their content was significantly lower than in *Gb* (Figure 13; Table S9).

**Figure 13.**
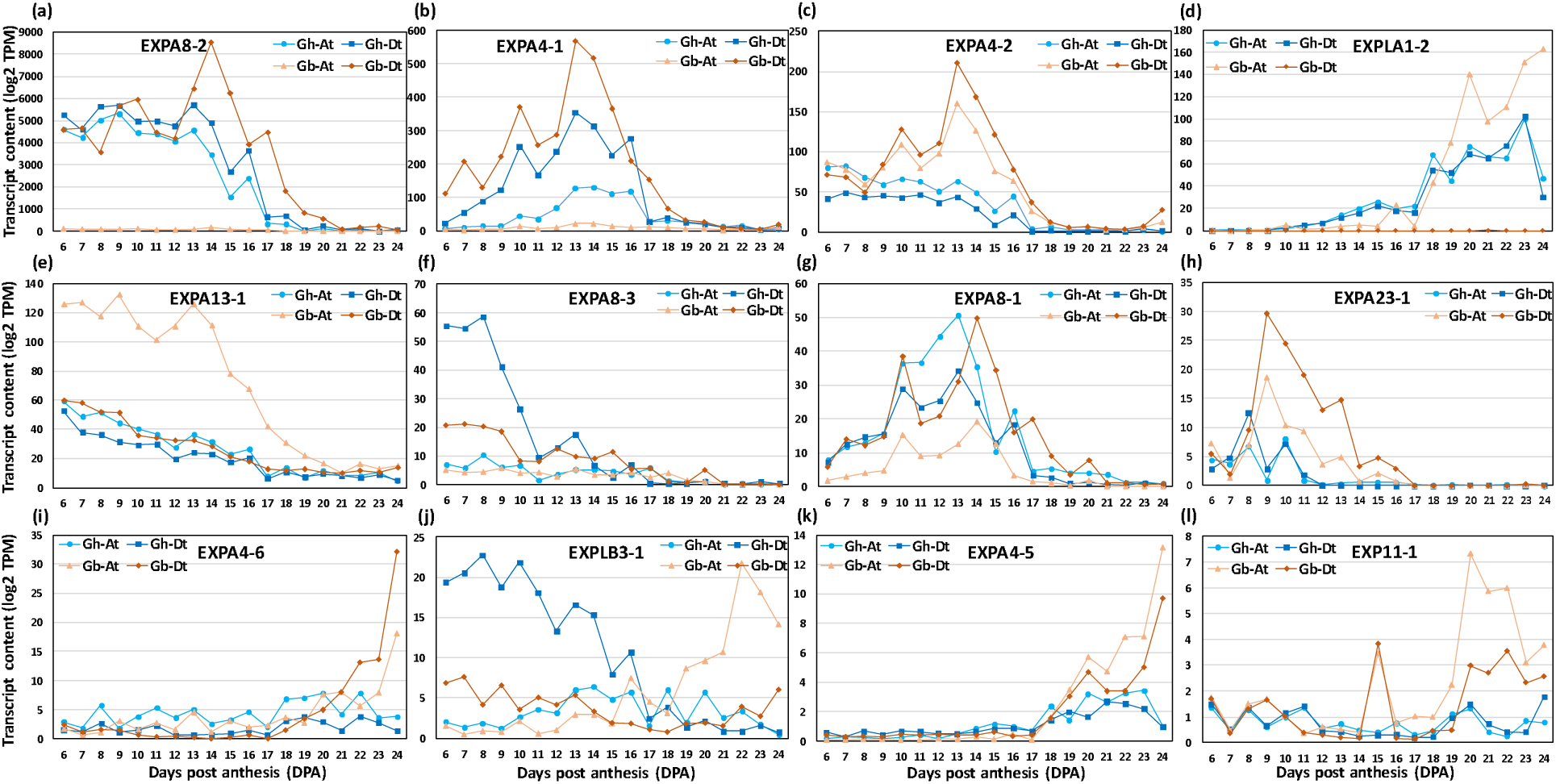
The profiles of expansin transcripts differentially expressed in G. hirsutum (Gh) and G. barbadense (Gb) fiber during development (6 - 24 DPA). Log2-transformed transcript per million (TPM) values of A sub-genome (At) and D sub-genome (Dt) of both the species are shown in the plot.

## DISCUSSION

*Gossypium hirsutum* (*Gh*) and *G. barbadense* (*Gb*) have different fiber properties with well-defined and overlapping stages of development. These species offer an excellent opportunity to study the dynamic temporal remodeling and development of a fiber cell that defines fiber characteristics, and comparative analysis between the species represents a natural form of experimental perturbation that can inform our understanding of the molecular underpinnings of the mature phenotypes. It is well known that *Gh* has moderate-quality fiber and that *Gb* offers superior quality (longer, stronger, and finer) fiber. Earlier studies pointed out that cotton fiber development includes tightly controlled complex gene expression networks, biosynthetic pathways, physiology, and development that eventually result in dynamic sequential changes in fiber CW polysaccharide epitopes, which decides the fiber quality in different species of cotton (Haigler et al., 2012; Jan et al., 2022; Jareczek et al., 2023). The main focus of our study was to analyze the fiber from both *Gh* and *Gb*, identify critical polysaccharide structures that drive fiber quality traits, and also identify potential glycosyltransferases that may be responsible for the synthesis of these polysaccharide structures.

The uniqueness and power of this large-scale comparative study is that glycome, transcriptome, and proteome profiling were conducted simultaneously on fibers from two different species, collected daily for 20 successive days (6 to 25 DPA), which were grown under controlled conditions. Our previous large scale glycome and transcriptome study on *Gh* fiber identified critical polysaccharides and several key putative glycosyltransferases that contribute to important fiber traits (Swaminathan et al., 2024). Although in this study proteomic analysis was conducted exclusively on *Gb* fiber, the integration of multi “omics” data demonstrated a broad correlation between the level of transcripts and corresponding glycosyltransferases and other polysaccharide-synthesizing enzymes detected in the membrane fraction in proteome (Figure 2).

This confirms the increased power of prediction of potential glycosyltransferases involved in the formation of specific polysaccharide structures/epitopes in the course of fiber development. Another important aspect of our study is that these 20 days were selected to cover the most critical stages of fiber development, comprising the fiber elongation, transition, and initial SCW thickening stages (Haigler et al., 2012). These stages of development are also characterized by high gene expression variation between *Gh* and *Gb*, which coincides with CW remodeling that potentially leads to differential fiber development between *Gh* and *Gb*, resulting in different fiber quality (Al-Ghazi et al., 2009; Liu et al., 2023). This comparative analysis, using coordinated analysis of transcriptome, proteome, and glycome, represents a promising approach to understand how the subtle details of cell wall polysaccharide alterations during key stages of cell wall synthesis might predict the underlying molecular machinery. Correlation analyses of polysaccharide compositional changes, expression levels of glycosyltransferase-encoding genes and their corresponding proteins at high temporal resolution allow us to better understand the dynamic changes that occur during fiber morphogenesis of the two most important commercial cotton species and identify their critical differences.

### Differences in cellulose accumulation between *G. hirsutum* (*Gh*) and *G. barbadense* (*Gb*) fibers

Here we observed a three-day delay in the rapid accumulation of cellulose content in *Gb* fibers compared to *Gh* during the transition period prior to SCW thickening (Figure 1). This observation agrees with the earlier studies regarding the extended fiber elongation phase reported at *Gb* in comparison to *Gh* (Tuttle et al., 2015; Hu et al., 2019; Pettolino et al., 2022). Such a delay in cellulose accumulation was reported previously even for different *Gb* accessions and different growing conditions (Lee et al., 2015; Pettolino et al., 2022). Coincidently, a similar delay was identified in the expression of the transcripts of ten SCW CESAs in *Gb* in comparison to *Gh* (Figure 8). In addition, transcripts of two of the PCW CESAs in *Gb* were expressed at a higher level for a longer period of time than in *Gh*. Previous transcriptomic studies utilizing only 3-time points (10, 15, 20 DPA) revealed only one of the PCW CESA (*CESA6*) having higher expression in *Gb* relative to *Gh* (Hernandez-Gomez et al., 2015). Several other studies that examined fewer time points of fibers also reported delayed and lower expression of some SCW CESAs in *Gb* relative to *Gh* (Lacape et al., 2012; Li et al., 2013; Hernandez-Gomez et al., 2015).

It was proposed that the delayed timing of cellulose accumulation might be one of the reasons for the superior fiber quality of *Gb* (longer, thinner, finer) relative to *Gh* (Lacape et al., 2012; Li et al., 2013; Pettolino et al., 2022). Also, it is well known that higher SCW CESA expression leads to a higher accumulation of cellulose at the SCW developmental stage, which results in thickening/rigidification of the CW, thus halting the CW elongation (Haigler et al., 2012). Our findings, combined with those of earlier reports, suggest that slower accumulation of cellulose (due to extended/higher PCW CESAs expression combined with delayed lower expression of SCW CESAs for a few days) results in delayed rigidification of CW, which might lead to longer fiber in *Gb*. Even though cellulose accumulation was delayed for about 3 days and was lower content in *Gb* at around 19 DPA compared to *Gh* (Figure 1), the cellulose accumulation in *Gb* increased rapidly later, and at the end of 25 DPA, both species had the same level of cellulose. Earlier, Li et al. (2013) reported that the CESA8 was the major player for SCW cellulose accumulation in both *Gh* and *Gb* fiber, and significantly contributed to rapid cellulose accumulation at later stages in *Gb* during SCW thickening stage. Interestingly, our transcriptome data also showed that the transcript level of only CESA8-B (Figure 8f) was high and equal in both species in comparison to all other SCW CESAs (Figure 8; Table S4).

#### Differences in matrix polysaccharides and differentially expressed glycosyltransferases that could contribute to differences in fiber quality between *G. hirsutum* (*Gh*) and *G. barbadense* (*Gb*)

Comparative analysis of the polysaccharide composition and related glycosyltransferases between *Gh* and *Gb* fiber revealed many interesting specifics that might be the cause for the fiber quality differences between these cotton species. First, it is worth pointing out the differences in the total amount of buffer-soluble and alkali-soluble polysaccharide fractions. In *Gh*, the content of buffer-soluble polysaccharides was higher than that of alkali-soluble at 6-7 DPAs, whereas in *Gb* these two fractions were present in more comparable amounts (Figure 1). However, the level of buffer-soluble polysaccharides reduced more rapidly in *Gh* than in *Gb* during development, and at later stages, the relative proportion of buffer-soluble polysaccharides remained at the same level in the two species. These differences are reflected in the differences in polysaccharide epitope distribution, discussed below.

Quantitative heat maps and SOM analyses of polysaccharide epitope profiles revealed significant differences between the two species, which guided a comparative analysis of the glycome and glycosyltransferase transcriptome data. The most interesting aspect of our finding was that some of the same polysaccharide epitopes from *Gh* and *Gb* fell into two different SOM groups, which suggests that polysaccharide distribution differs in fiber CW between the two species. Although some of these particular differences seem not substantial and are statistically insignificant, even subtle differences, taken together, could potentially impact fiber growth and development. In particular, the polysaccharide epitopes from our third categorical group (Figure 7) may be a promising target for further detailed investigations.

Our large-scale glycome profiling showed that most of the buffer and alkali-extracted fucosylated and non-fucosylated xyloglucan epitopes were very low in *Gb* in comparison with *Gh* (Figures 6 and 7), which agrees with similar earlier comparative studies (Avci et al., 2013; Hernandez-Gomez et al., 2017; Guo et al., 2019). These earlier studies reported that the presence of xyloglucan was lower in cotton fiber middle lamellae (CFML) and PCW from *Gb* than in *Gh*. The authors proposed that the role of CFML is to adhere the fibers in bundles to promote fiber elongation. However, Avci et al. (2013) reported that CFML is not required for fiber elongation in *Gb*, since it continues to elongate rapidly even after lysis of CFML. Additional studies will be required to confirm the correlation between the amount of xyloglucan and the dynamics of CFML/PCW development in promoting fiber length in different species of cotton.

Interestingly, in our transcriptome profiling study, we found that the transcript levels of all GTs related to xyloglucan biosynthesis (*CSLCs/XXTs/MUR3s/XLT*) (Figure 9; Table S4) were lower in *Gb* than in *Gh*, which may be related to the lower xyloglucan epitope content in *Gb* than in *Gh*. A previous study by Guo et al. (2019) using fiber at 10, 15, and 20 DPA showed that the transcript levels of a *CSLC4* and a *MUR3* were lower in *Gb* than in *Gh*. In our transcriptome data, we found a significantly larger set of various xyloglucan-synthesizing glycosyltransferases from both A and D genomes that were expressed at a lower level in *Gb* in comparison to *Gh*. It will be interesting in future studies to examine the contribution of all these enzymes to the superior quality fiber in *Gb*.

It is well known that xyloglucans are highly cross-linked with cellulose microfibrils through non-covalent hydrogen bonds to form a “tethered network” that gives rigidity to the CW and restrains cell expansion (McCann et al., 1990; Somerville et al., 2004; Park & Cosgrove, 2015). Interestingly, some of the earlier studies showed that the short fiber mutants (Ligon lintless-1 & 2; Li_1_/ Li_2_), had a higher accumulation of xyloglucan content during elongation stages due to higher expression of xyloglucan-synthesizing glycosyltransferases, which resulted in an extreme reduction in fiber length (Shao et al., 2011; Naoumkina et al. 2017). Overexpression of a MUR3 (GhMUR3-2Dt; Gohir.D10G136900) in *Gh* cotton resulted in shorter and thicker fiber compared to wild type (Wu et al., 2024). All data accumulated so far suggest that a higher amount of xyloglucan negatively correlates with fiber length, most likely impacting the “tethered xyloglucan-cellulose network”, which leads to an increase in the rigidity of fiber CWs. Thus, the results suggest that a lower amount of xyloglucan seems to loosen this network which helps to extend elongation time and promotes longer fiber in *Gb* in comparison with *Gh*.

Another hemicellulosic polysaccharide, xylan, is the major component in SCWs, and due to its interaction with cellulose and lignin, xylans are essential for SCW architecture and strength (Gille & Pauly, 2012). In our previous study, we showed that some of the xylan epitopes (Xyl-3Ar, Xyl-MeGlcA) and associated glycosyltransferases rapidly peak at the transition stage in *Gh* fiber before the rapid SCW cellulose accumulation/CW thickening and before the changes in microfibril orientation (Swaminathan et al., 2024). Here, we found that the amount of loosely bound buffer-extractable xylans, mostly represented by the xylan backbone epitopes (Xyl-BB) and a glucuronoxylan epitope (Xyl-GlcA), was lower in *Gb* than in *Gh*. The alkali-extractable arabinoxylans (Xyl-2Ar, Xyl-3Ar) and a methylated glucuronoxylan (Xyl-MeGlcA-2) showed different profiles between *Gh* and *Gb* (Figure 7), and Xyl-MeGlcA-2 present in higher amounts in *Gb* than in *Gh*. Somewhat similar differences in xylan-related epitope profiles were reported earlier (Avci et al., 2013; Hernandez-Gomez et al., 2015).

We compared the expression levels of xylan-synthesizing glycosyltransferases between the two species. Transcripts of most of the *IRXs/FRA8/PARVUSs/ESKs/TBLs/RWAs* genes, which are involved in xylan backbone synthesis, were expressed more highly in *Gh* than in *Gb*, which correlates well with the presence of higher amounts of xylan backbone epitopes in *Gh* than in *Gb* (Figure 10). On the other hand, some of the *GUXs* and *GXMTs*, which are responsible for the synthesis of methylated glucuronoxylans, were expressed more highly in *Gb* than in *Gh*, which correlates with the higher content of Xyl-MeGlcA-2 epitopes detected in *Gb* (Figure 7).

Recent reverse-genetic studies in cotton using some of the xylan-synthesizing enzymes (FRA8/IRX9/10/14/15) from *Gh* showed that xylan synthesis positively correlates with cellulose content, cellulose microfibril orientation, SCW cellulose deposition, fiber length, and thickness (Li et al., 2014; Chen et al., 2020a; Guo et al., 2024; Li et al., 2024). In our previous large-scale profiling study (Swaminathan et al., 2024), we also observed that three xylan epitopes were highly correlated with the cellulose content, the expression of SCW CESAs, the cellulosic microfibril orientation and the CW thickness phenotype of *Gh* fiber. The function of heteroxylans (glucuronoxylans and arabinoxylans) is usually related to the strengthening of CW by acting as a guiding scaffold for cellulose microfibril orientation/arrangement (Grantham et al., 2017; Smith et al., 2017; Crowe et al., 2021; Pfaff et al., 2024). The strength of the cotton fiber CW is an important quality essential for textile industry processing. The differential profiles and contents of methylated glucuronoxylan (Xyl-MeGlcA) and arabinoxylans (Xyl-2Ar and Xyl-3Ar) observed in *Gb* could potentially contribute to its longer, stronger and finer quality fiber. In our proteome data, we also noticed that several IRXs, GUXs, GXMTs, XATs are abundantly present and matched their corresponding transcript profiles (Figure 2; Table S6). Additional research is still required to understand the reported differences in heteroxylan accumulation in fiber and its impact on fiber quality. Interestingly, an earlier study of *Arabidopsis* mutants showing various levels of glucuronoxylan deficiency demonstrated that the level of glucuronoxylans is critical for the precise cellulose network formation and CW integrity of SCWs (Crowe et al., 2021). This might suggest that temporal synchrony in the presence of appropriate heteroxylans quality and quantity in combination with the other polysaccharides, affects the CW architecture and, thus, the final quality of the fiber.

Pectins are the major components of the PCW and CFML (up to 35% dry weight) of the cotton fiber (Kaczmarska et al., 2022). There are four main structural components of pectins: homogalacturonans (HGs), Rhamnogalacturonans-I (RG-I), RG-II, and xylogalacturonans. The cellulose and hemicellulose networks are embedded in a matrix of pectins and proteins in PCW. HGs and RGs play a central role in regulating the viscosity/extensibility/gelling property of the CW matrix, thus controlling polysaccharide interactions, CW elongation, and cell growth (Hwang & Kokini, 1992; Yapo, 2011; Sousa et al., 2015; Zheng et al., 2020). Our results from glycome profiling showed that a methyl-esterified HG epitope (HG-BBMe-K) and most of the highly branched RG-I epitopes were lower in *Gb* relative to *Gh*. In contrast, the de-esterified HGs (HG-BBde), and RG-I backbone epitopes (RG-I-BB, Gal4-BB) were present in equal amounts in both species. Similar observations were reported earlier by Avci et al. (2013). Another comparative “omics” study (Liu et al., 2013) reported the presence of lower amount of methyl-esterified HG in *Gb* compared to *Gh* during elongation time and proposed that pectin content might be one of the important factors influencing the fiber elongation and quality. Our results suggest that the presence of a lower content of HGs and highly branched RG-Is during the PCW developmental/fiber elongation stage in *Gb* could support its longer fiber phenotype.

Our comparative transcript analysis showed that most of the glycosyltransferases that synthesize the HG epitopes, including methyl and acetyltransferases (GAUTs, GATLs, CGRs/QUAs, TBLs/TBRs) were present in higher amounts in *Gh* than in *Gb*. A similar trend was observed for RG-I-synthesizing glycosyltransferases, most likely resulting in a higher amount of pectin epitopes in *Gh* than in *Gb*. The cotton AG:GALT (Gohir.A04G041100/*Gh*AG:GALT1-2At) protein was found to play a functional role in galactosylation of arabinogalactan chains of RG-I/AGPs molecules (Qin et al., 2017). The *AG:GALT1-2At* RNAi silenced cotton plants, had less galactose content in the RG-I and produced longer fibers as a result of CW loosening. Conversely, AG:GALT1-2At-overexpressing cotton plants had the opposite effect. All these results indicate that higher amounts of pectin epitopes during the elongation stage lead to broader hydrogen bonding interactions between the side chains of pectin molecules, which causes gelling and increased firmness of CW, which could negatively impact the elongation time and length of cotton fibers. A more detailed comparative investigation of glycosyl hydrolases contributing to cotton fiber development will be our next effort.

### Differentially expressed transcripts of expansins of *G. hirsutum* (*Gh*) and *G. barbadense* (*Gb*) fibers

Expansins are CW-loosening protein that act in a non-enzymatic manner to weaken the noncovalent bonds in hemicellulose-cellulose networks, resulting in slippage of cellulose microfibrils, and ultimately leading to cell expansion (Sampedro & Cosgrove, 2005). The importance of expansins, as one of the major factors in cotton fiber elongation, was reported earlier (Xu et al., 2013; Li et al., 2016; Lv et al., 2020). Therefore here, in addition to glycosyltransferases, we examined the transcript levels of reported expansins in both *Gh* and *Gb* fibers. Interestingly, the transcript level of most of the highly expressed expansins was higher in *Gb* relative to *Gh* (Figure 13; Table S9). Additionally, the expression of some of the expansins was significantly higher in *Gb* through the later stages (after 17 DPA), which coincides with the extended elongation stage in this cotton genotype. In contrast, in *Gh*, the transcript level of most of the expansins decreased at 17 DPA, coinciding with the transition stage before the SCW thickening and complete cessation of fiber elongation in this genotype. Our results suggest that the temporally controlled higher level of expansin expression plays an important role in extended fiber elongation and the development of superior fiber quality in *Gb*.

Previous reverse-genetic studies of EXPA8-2 (the highest expressing expansin from both *Gh* and *Gb*) by Xu et al. (2013) and Li et al. (2016) showed that the cotton fiber length is increased in *Gb*EXPA8-2 overexpressing lines and reduced in the silenced lines. Overexpression of GbEXPA8-2 also altered the SCW-associated genes and delayed the accumulation of SCW cellulose, which resulted in extended elongation time of the PCW stage and long fibers in *Gb*. A separate transgenic experiment in *Gh* by overexpressing *Gh*EXPA8-2 along with BURP-domain containing protein resulted in the increased fiber length (Xu et al., 2013).

In summary, this study integrated large-scale CW polysaccharide glycome profiling and glycosyltransferase transcriptome profiling of the two commercially important cotton species*, G. barbadense and G. hirsutum,* which have different fiber qualities. Our comparative study at high temporal resolution discovered potentially critical polysaccharide epitopes and polysaccharide-synthesizing glycosyltransferases that may determine or contribute to the differences in final fiber quality and characteristics in cotton. The presence of relatively lower amounts of methyl esterified HG, highly branched RG-I epitopes, xyloglucan, and xylan epitopes most likely impact both the length and strength of cotton fiber by extending fiber elongation time in *G. barbadense* in comparison to *G. hirsutum*. Higher and prolonged expression of PCW CESAs, delayed expression of SCW CESAs, and corresponding delayed cellulose accumulation are the most plausible factors contributing to the longer fiber phenotype of *G. barbadense*. Additionally, our study suggests that the differentially expressed heteroxylans (Xyl-2Ar, Xyl-3Ar and Xyl-MeGlcA), might also be important for cellulose microfibril arrangement and different strengths of the fibers in both species. Our study, together with multiple earlier reports, pointing that a particular polysaccharide structures and levels of glycosyltransferases synthesizing them might contribute to the differences in fiber quality. Recently, the prospects of using molecular genetics to engineer cotton fiber phenotypes have shown promise. Therefore, the newly obtained insights into the molecular control of fiber morphology and quality will be a valuable source of information for future steps in not only refining the specific factors dictating the final quality of cotton fiber but also providing directions for quality improvement through cotton transformation.

### EXPERIMENTAL PROCEDURES

#### Cotton plant growing conditions and boll collection

Two different cotton species, *Gossypium hirsutum* (*Gh*; accession TM-1) and *G. barbadense* (*Gb*; accession 3-79) were grown synchronously under controlled conditions in a growth chamber (Conviron E-15, Controlled Environments Inc. ND, USA). Plants were grown individually in a two-gallon pot containing a potting mix (4:2:2:1 ratio of soil:perlite:bark:chicken grit). Growth chambers were set for 16-hour days with temperature of 28° C and 500 µmol of light. Flowers were self-pollinated before noon every day and tagged. For 20 different time points (from 6 to 25 DPA), bolls were collected at mid-day, and stored at −80°C. For each time point, three biological replicate bolls were used for glycome, proteome and transcriptome analysis.

#### Cell wall and polysaccharide extraction

From the cotton fiber, the cell wall (CW) and the buffer-soluble and alkali-soluble polysaccharide fractions were extracted as per the established protocol (Avci et al., 2013; Swaminathan et al., 2024). In brief, using a razor blade and a tweezer, fibers were isolated from the developing seeds and adequate care was taken to avoid the seed coat while harvesting the fibers. Harvested cotton fibers from a single boll were ground into powder using liquid nitrogen, and the CW was then extracted from the powdered fiber using solvents (Swaminathan et al., 2024). Polysaccharides fractions were sequentially extracted from the CW by first using 50mM CDTA:50mM ammonium oxalate (1:1) buffer, followed by 4M KOH to extract buffer-soluble and alkali-soluble polysaccharides, respectively (Avci et al., 2013; Swaminathan et al., 2024). The final pellet, which contained a mixture of amorphous and crystalline celluloses, was weighed. The crystalline cellulose content present in the final pellet was measured by using the Updegraff reagent (acetic acid: nitric acid: water, 8:1:2 v/v) (Updegraff, 1969).

#### Glycome profiling of epitopes of buffer-soluble and alkali-soluble polysaccharide fractions

Glycome profiling was carried out using a standard protocol (Pattathil et al., 2012). Initially, the sugar content in each of the buffer-soluble and alkali-soluble polysaccharide fractions from each sample was estimated by the phenol-sulfuric acid method (Pattathil et al., 2012). Later, the polysaccharide sample was dissolved in water and an equal amount of 3 µg (50 µl/well from a 60 µg/µl solution) was added in each well in 96-well ELISA plates (Costar 3595) and the polysaccharides were allowed to bind to the bottom of the well by drying in an oven at 37°C. Glycome profiling was conducted using an ELISA method with 71 different polysaccharide epitope-specific antibodies (Table S2), as described earlier (Pattathil et al., 2010; Swaminathan et al., 2024), and these antibodies were selected based on the literature (Avci et al., 2013; Thorne et al., 2023). Data from the ELISA for each epitope were obtained from either buffer-soluble or alkali-soluble polysaccharide fractions of the same sample (Table S4). To distinguish between the two types of polysaccharide fractions, for the naming of epitopes, we used a naming convention of where a suffix of “-C” refers to the buffer-soluble fraction (50mM CDTA:50mM ammonium oxalate buffer extracts) and “-K” refers to the alkali-soluble fraction (4M KOH extracts).

#### Self-organizing map (SOM) clustering

The Self-Organizing Map (SOM) is a robust machine-learning method used for clustering analysis (Kohonen, 1990). An SOM algorithm, based on an inner product distance metric, was applied to the ELISA-based glycome profiling results to cluster the abundance profiles of glycome epitopes from two separate interpolated datasets: buffer-soluble and alkali-soluble polysaccharide fractions from two different species of cotton (Wehrens & Kruisselbrink, 2018). The SOM grid structures consisted of 3 rows and 4 columns for both buffer-soluble and alkali-soluble polysaccharide fractions datasets. In the SOM analysis, the data from both the species for each of the fractions were combined, and two different colors were used to maintain the identity of polysaccharide epitopes.

#### RNA extraction and RNA sequencing

In parallel to glycome profiling, RNAseq and transcriptomic profiling were carried out for the cotton fibers collected simultaneously for the same days (6 to 25 DPA) (Grover et al., 2025). Briefly, a modified version of the Sigma Plant Spectrum Total RNA kit (Sigma-Aldrich) was used to extract RNA from the fibers. Library construction (NEBNext Ultra II RNA Library Prep Kit) and sequencing (as PE150; Illumina NovaSeq 6000) were conducted by the Iowa State University DNA facility. Data were processed using Trimmomatic version 0.39 (trimmomatic/0.39-da5npsr) (Bolger et al., 2014) from Spack (Gamblin et al., 2015) for read and quality trimming. The genes from transcriptome were annotated based on the published reference *Gh* genome (Chen et al., 2020b) and *Gb* genome (Chen et al., 2020b). The transcripts per million (TPM) output by Kallisto for each sample were imported into R/4.2.2 (R Core Team 2022) and RNA-seq quality was assessed by consistency of number of genes expressed over time and among replicates. The number of expressed genes per sample (TPM > 0) was plotted across developmental time using ggplot2, and visual outliers were discarded.

#### Transcriptomic analysis of glycosyltransferases

Glycosyltransferases and enzymes that decorate polysaccharides play key roles in the synthesis of fiber CW polysaccharides. To compare the expression of these enzymes between the two species, first, we first compiled a list ofknown or putative Arabidopsis glycosyltransferases based on our previous study and other studies (Table S7). Subsequently, the gene sequences of *Arabidopsis* proteins were used to find out the homologous genes of *Gh* and *Gb* from the Phytozome 13 (https://phytozome-next.jgi.doe.gov) and the Cotton Functional Genomics Database (https://cottonfgd.net). Based on this information, expression profiles of glycosyltransferases and other polysaccharide-decorating enzymes in both *Gh* and *Gb* were generated from the transcriptome data (Grover et al., 2025). Comparisons of expression profiles of transcripts of these enzymes between *Gh* and *Gb* and their sub-genomes (At and Dt) were conducted.

#### Microsomal protein profiling

The microsomal protein fraction (P200) was obtained from intact cotton fiber tissue at 6 to 25 DPA, as described previously (Lee et al., 2025). Briefly, apoplastic proteins and extracellular vesicles were removed from the intact ovules by dipping each intact ovule into 5 mL of pre-chilled microsome isolation buffer (MIB) for 10 minutes. Subsequently, the fiber tissues were isolated, homogenized in cold MIB using a Polytron homogenizer and filtered through cheesecloth. From the filtrate, microsomes were enriched using an ultracentrifuge (200,000 x g for 20 minutes at 4 °C). The final pellet was mixed with 8M Urea to denature membrane proteins, digested with trypsin, column purified and the peptides were analyzed using Bruker’s TIMS-TOF HT (Bruker Daltonics GmbH) mass spectrometer (MS), which was coupled with a reverse-phase liquid chromatography system, nanoElute2 (Bruker Daltonics GmbH). The MS raw data were processed using the directDIA™ approach in Spectronaut software (Version 19.4, Biognosys) following the vendor’s recommendations. The spectral library was constructed using a cotton FASTA file containing 55,237 entries and DPA-pooled DDA runs. The false discovery rate (FDR) was set to 1% at both the peptide and protein levels. The glycosyl transferase proteomic data presented here (Table S5) are part of a larger dataset that is in preparation for publication.

#### Statistical analysis

The experimental data, including Pearson correlation coefficient (PCC) analysis were subjected to statistical analysis using R software (R Studio; R Core Team, 2022). Data from the corresponding polysaccharide epitopes from each of the two-cotton species were initially tested for significance using analysis of variance (ANOVA). Subsequently, Fisher’s protected least significant difference (LSD) test was used to compare the means (at *p* < 0.05). Based on the significance differences, the epitopes were grouped into mainly three different categories.

## ACKNOWLEDGEMENTS

This study was supported by the NSF PGRP Award No. 1951819. We thank Dr. Uma Aryal and Dr. Venkatesh Thirumalaikumar at the Purdue Proteomics Facility for running the LC/MS samples.

## AUTHOR CONTRIBUTIONS

DBZ, JFW, OAZ, and JX conceived the research. SS, YL, and CEG grew the cotton plants, pollinated, collected bolls. SS, ASM, LES, and MFD extracted polysaccharides from cotton fibers. CEG extracted RNA from fibers. CEG and JFW analyzed the RNAseq transcriptome data. SS, and OAZ conducted glycome profiling and analyzed the data. PY, JX, and MFD generated SOM. YL, and DBS extracted proteins, conducted proteome experiments and analyzed the data. SS wrote the manuscript, with significant input from OAZ, JFW, and DBS. All authors contributed with comments and suggestions to finalize the manuscript.

## DATA AVAILABILITY STATEMENT

RNAseq reads of cotton fibers collected from 6 to 25 DPA were deposited into the NCBI-SRA under PRJNA1099209 (*Gossypium hirsutum*) and PRJNA1222456 (*Gossypium barbadense*). All the data and materials that support the findings of this study are available upon request from the corresponding author.

## SUPPORTING INFORMATION

**Figure S1.** Pictorial representation of structures of analyzed cell wall (CW) polysaccharides epitopes and glycosyltransferase enzymes involved in their synthesis.

**Table S1.** Fiber cell wall polysaccharides of Gossypium barbadense and Gossypium hirsutum (6 - 25 DPA).

**Table S2.** List of polysaccharide specific antibodies used in this study and their binding epitopes.

**Table S3.** Glycome profiling data of Gossypium barbadense and Gossypium hirsutum (6 - 25 DPA).

**Table S4.** List of cell wall polysaccharide synthesizing enzymes from Arabidopsis and cotton.

**Table S5.** Proteome profiling data of glycosyltransferases.

**Table S6.** Pearson Correlation Coefficient (PCC) analysis of Gossypium barbadense glycome, proteome and transcriptome profiles data.

**Table S7.** Self-organizing map (SOM) groups of glycome profiled polysaccharide fractions and the epitopes present in each group.

**Table S8.** Categories of polysaccharides epitopes of Gossypium barbadense and Gossypium hirsutum (6 - 25 DPA) based on SOM.

**Table S9.** List of cotton expansins genes homologous to genes of Arabidopsis expansins.

