## Supplemental Figure 1 for "Comparative multi “omics” profiling of *Gossypium hirsutum* and *Gossypium barbadense* fibers at high temporal resolution reveals key differences in polysaccharide composition and associated glycosyltransferases"

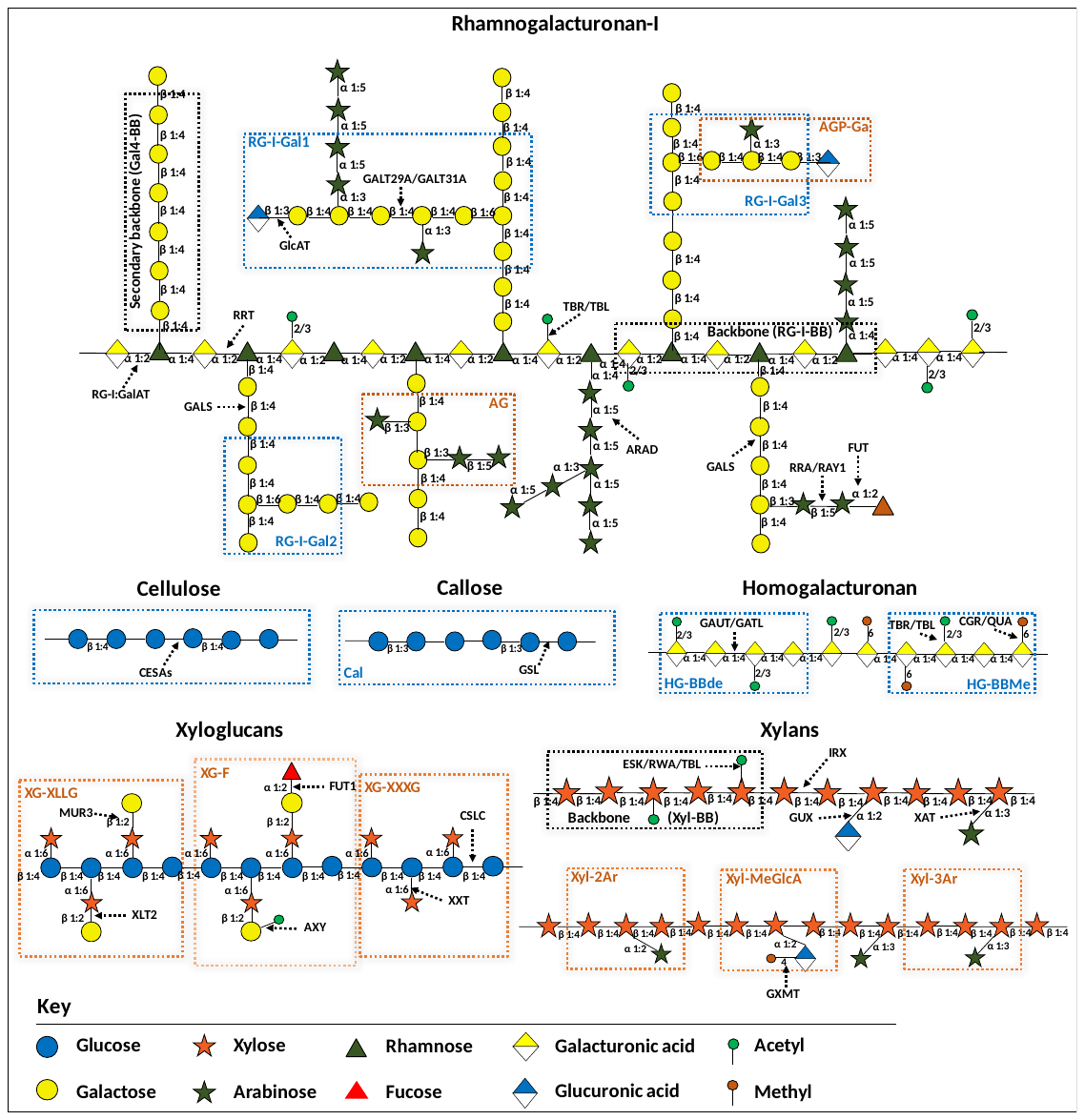

**Figure S1.** Pictorial representation of structures of analyzed cell wall (CW) polysaccharides epitopes and glycosyltransferase enzymes involved in their synthesis. The schematic structure of Rhamnogalacturonan-I (RG-I) pectic polysaccharide displays different epitopes like, backbones (RG-I-BB, Gal4-BB), β-1,6-linked galactans (RG-I-Gal1, RG-I-Gal2 and RG-I-Gal3), and arabinogalactans (AG, AGP-Ga). Cellulose and callose polysaccharides are made up of glucose molecules connected by β-1,4- and β-1,3-linkages, respectively. Homogalacturonan (HG) pectic polysaccharide displays two different epitopes, Methyl-esterified HG (HG-BBMe), and de-esterified HG (HG-BBde). Xyloglucan (XG) structure shows, xylosylated XG (XG-XXXG), galactosylated XG (XG-XLLG), and fucosylated XG (XG-F) epitopes. The schematic structures of xylans show the backbone (Xyl-BB), arabinoxylans (Xyl-2Ar and Xyl-3Ar) and methylated-glucuronoxyaln (Xyl-MeGlcA) epitopes. Refer to Table S2 for the details of the epitopes recognized by the antibodies used in this study. The known or putative Arabidopsis CW synthesizing glycosyltransferase enzymes are denoted by black dotted arrows.
